# A single bacterial sulfatase is required for metabolism of colonic mucin *O*-glycans and intestinal colonization by a symbiotic human gut bacterium

**DOI:** 10.1101/2020.11.20.392076

**Authors:** Ana S. Luis, Chunsheng Jin, Gabriel Vasconcelos Pereira, Robert W. P. Glowacki, Sadie Gugel, Shaleni Singh, Dominic P. Byrne, Nicholas Pudlo, James A London, Arnaud Baslé, Mark Reihill, Stefan Oscarson, Patrick A. Eyers, Czjzek Mirjam, Gurvan Michel, Tristan Barbeyron, Edwin A Yates, Gunnar C. Hansson, Niclas G. Karlsson, Alan Cartmell, Eric C. Martens

## Abstract

Humans have co-evolved with a dense community of microbial symbionts that inhabit the lower intestine. In the colon, secreted mucus creates a physical barrier that separates these microbes from the intestinal epithelium. Some gut bacteria are able to utilize mucin glycoproteins, the main mucus component, as a nutrient source. However, it remains unclear which bacterial enzymes initiate the degradation of the highly complex *O*-glycans found in mucins. In the colon, these glycans are heavily sulfated, but the specific sulfatases that are active on colonic mucins have not been identified. Here, we show that sulfatases are essential to the utilization of colonic mucin *O*-glycans by the human gut symbiont *Bacteroides thetaiotaomicron*. We have characterized the activity of 12 different sulfatases encoded by this species, showing that these enzymes collectively are active on all of the known sulfate linkages in colonic *O*-glycans. Crystal structures of 3 enzymes provide mechanistic insight into the molecular basis of substrate-specificity. Unexpectedly, we found that a single sulfatase is essential for utilization of sulfated *O*-glycans *in vitro* and also plays a major role *in vivo*. Our results provide insight into the mechanisms of mucin degradation by gut bacteria, an important process for both normal microbial gut colonization and diseases such as inflammatory bowel disease (IBD). Sulfatase activity is likely to be a keystone step in bacterial mucin degradation and inhibition of these enzymes may therefore represent a viable therapeutic path for treatment of IBD and other diseases.

## Introduction

The human gut microbiota (HGM) significantly impacts several aspects of intestinal health and disease, including inflammatory bowel disease (IBD)^1^ and colorectal cancer (CRC)^2^. In the colon, secreted mucus creates a physical barrier that separates gut microbes from the intestinal epithelium^3^ preventing close contact that can lead to inflammation and eventual CRC if this barrier is either experimentally eliminated^4, 5^ or has reduced glycosylation^6–9^. A major component of the colonic mucus is mucin 2 (MUC2), a glycoprotein that contains up to 80% glycans by mass and more than 100 different glycan structures that are *O*-linked to serine or threonine residues^10^. Mucin glycosylation is variable along the gastrointestinal (GI) tract with a marked increase in sulfation in the colon^11^. In mucins, *O*-linked sulfate may be attached to the 6-hydroxyl of *N*-acetyl-D-glucosamine (6S-GlcNAc) and non-reducing end D-galactose (Gal) sugars at hydroxyl positions 3-, 4- or 6- (3S-, 4S- and 6S-Gal, respectively)^11–13^ (**Fig. 1a**). Sulfation often occurs as terminal caps that block enzymatic degradation of oligosaccharides. To degrade and utilize colonic mucin *O*-glycans, members of the HGM need to express appropriate carbohydrate sulfatases to remove these modifications. *Bacteroides thetaiotaomicron* (*Bt*) is a dominant member of the human gut microbiota that is able to utilize *O*-glycans as a sole nutrient source^14^. Underscoring the importance of sulfatases, *Bt* requires active sulfatases for competitive colonization of the wild-type mouse gut^15^ and to induce inflammation in genetically-susceptible mice^16^. However, the specific sulfatases that mediate these effects remain unknown. Indeed, despite the critical roles of sulfatases in many biological processes, including several human diseases^17^, a significant knowledge gap exists regarding the biochemical, structural and functional roles of these enzymes.

**Figure 1.**
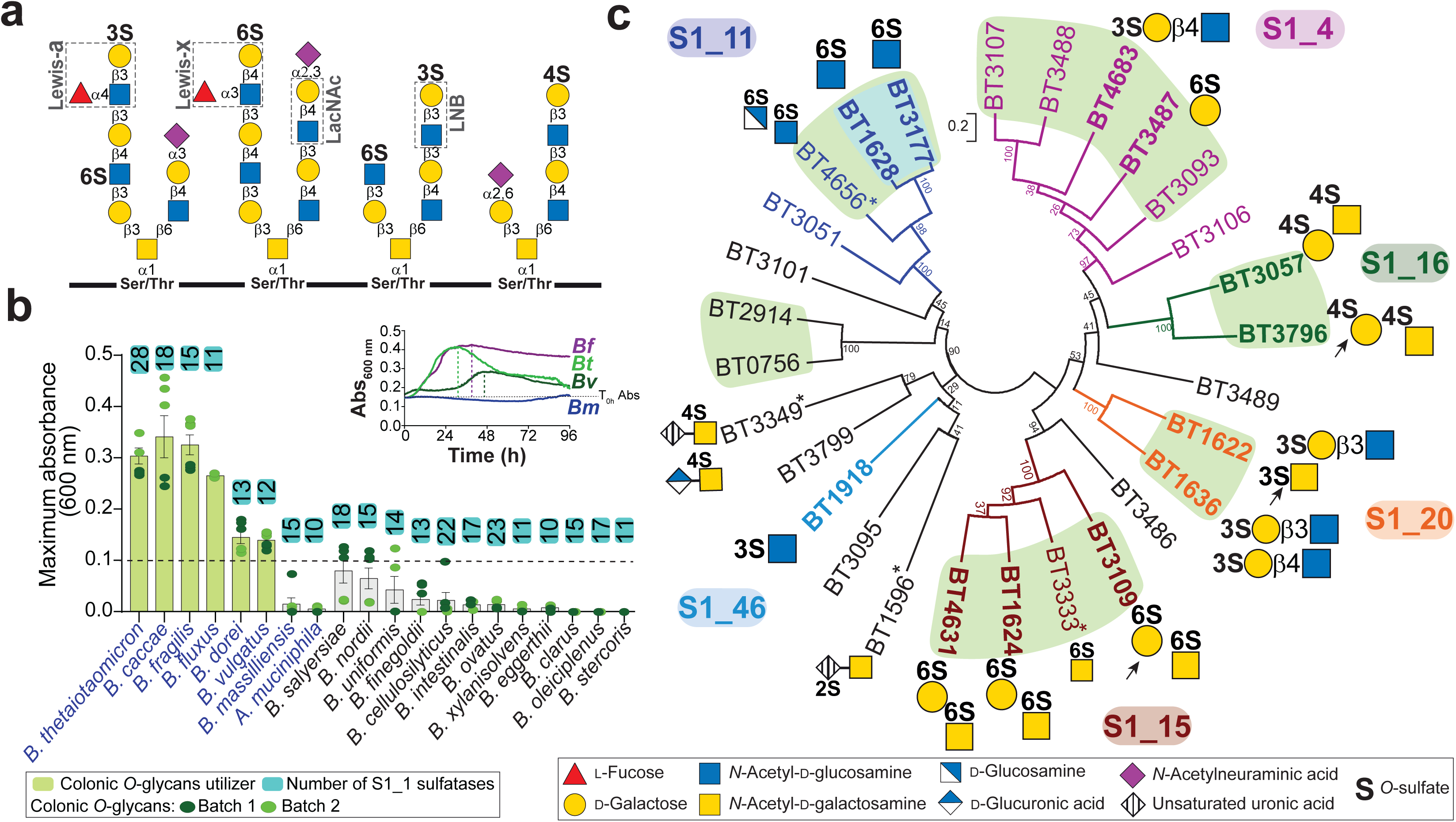
Bacterial growth on colonic mucin and *Bt* sulfatase activities. **a,** Schematic representation of mucin *O*-glycans and relevant terminal epitopes (dashed boxes). Sugars are shown according to the Symbol Nomenclature for Glycan system^60^. **b**, Growth of *Bacteroides* type strains and A*kkermansia muciniphila* on colonic mucin *O*-glycans (cMO) and number of respective encoded S1 sulfatases. The bars represent the average of two independent experiments with different batches of cMO. Bacterial species able to utilize gastric mucin glycans are highlighted in blue. Maximum absorbance is the difference of the maximum absorbance value (Abs_600nm_) for each culture and the initial absorbance at time 0 (T_0h_). Graphic shows the example of growth curves for *B. fragilis* (*Bf*), *B. thetaiotaomicron* (*Bt*), *B. vulgatus* (*Bv*) and *B. massilliensis* (*Bm*). **c**, Phylogeny of *Bt* sulfatases showing the 28 S1 sulfatases and their respective substrates where known, including this study. Enzymes are color coded according the respective subfamilies with sulfatases characterized in this study highlighted in bold. * indicates sulfatase activity previously characterized and arrows point the substrate preferentially targeted by the respective enzyme. Sulfatases on a shared branch that share more than 86% and 39-58% of sequence identity are highlighted in blue and green background, respectively. Data from biological replicates n = 3 to 6 and error bars denote s.e.m..

We hypothesized that specific *Bt* sulfatases play essential roles in initiation of *O*-glycan degradation. To test this, we measured the activities of 23 putative *Bt* sulfatases, determining that 12 enzymes are active on either model glycan substrates or purified colonic mucin *O*-glycans. Together with defining the specific activities of these enzymes, we determined the corresponding structures for 3 sulfatases, revealing the basis of substrate specificity. Using molecular genetics, we next assessed the contributions of these sulfatases to *Bt* fitness *in vitro* and *in vivo,* unexpectedly revealing that a single enzyme is essential for utilization of sulfated mucin *O*-glycans and plays a major role in competitive gut colonization. Identifying specific bacterial sulfatases that are critical for intestinal mucin degradation provides new potential targets, with a goal of blocking progression of diseases such as IBD and possibly other disorders that result from bacterial disruption of the mucus barrier.

### Utilization of colonic mucins by HGM species

Several studies have identified HGM members that are able to utilize porcine gastric mucin *O*-glycans (gMO) as a sole carbon source^14, 18, 19^. However, this substrate does not adequately reflect the structural complexity of mucin *O*-glycans found in the colon, especially those with increased sulfation that are lacking in gMO^11^. To identify HGM species that utilize sulfated colonic mucins, we measured the growth of 19 *Bacteroides* type strains, plus *Akkermansia muciniphil*a, on highly sulfated porcine colonic mucin oligosaccharides (cMO). We identified six *Bacteroides* strains that utilize cMO (**Fig. 1b, Extended Data Fig. 1**). Interestingly, two known mucin-degraders, *A. muciniphila*^20^ and *B. massiliensis*^21^, grew robustly on gMO but failed to utilize sulfated cMO as a substrate (**Extended Data Fig. 1**), highlighting the importance of employing colonic mucins as substrates to draw more physiologically-relevant conclusions about the metabolic targets of these organisms. *Bt*, the bacterium with the highest number of sulfatases (28), was one of the strains with the best growth on cMO (**Fig. 1b**), suggesting that some of these enzymes might play key roles in promoting this ability. Therefore, to understand the role of sulfatases in colonic mucin utilization by HGM bacteria we focused on the biochemical and genetic characterization of the *Bt* enzymes. See **Supplementary Discussion 1** for additional details of GI mucins and bacterial growth kinetics.

### Substrate specificity of Bt sulfatases

Sulfatases are classified into four main families (S1 to S4) in the SulfAtlas database according to sequence similarity, catalytic mechanism and fold^22^. Family S1 is currently divided into 72 subfamilies (designated S1_X) and comprises the formylglycine sulfatases, which operate via a hydrolytic mechanism that utilizes a non-genetically coded formylglycine amino acid as its catalytic residue. In *Bt* and other anaerobic bacteria, this residue is introduced co-translationally by the anaerobic sulfatase maturating enzyme (anSME)^15^, which converts a serine or cysteine, within the consensus sequence **C/S**-X-P/A/S-X-R, to formylglycine, which then serves as catalytic nucleophile^23^. The *Bt* genome encodes 28 S1 sulfatases classified into twelve different subfamilies (**Supplementary Table 1**). Four *Bt* sulfatases have been previously characterized and all act on glycosaminoglycans (GAGs) that are components of extracellular matrix (**Fig. 1c**)^24, 25^. Interestingly, several of the uncharacterized S1 sulfatases are encoded within polysaccharide utilization loci (PULs) that are known to be upregulated *in vivo* or during growth on gMO^14^ and encode other glycoside hydrolases enzymes potentially involved in degrading mucin *O*-glycans (**Extended Data Fig. 2**).

To understand the role of sulfatases in mucin metabolism, we cloned and expressed in soluble form 23 of the remaining 24 uncharacterized sulfatases. The recombinant proteins were tested for activity against a panel of commercially available sulfated saccharides (**Supplementary Table 2**). In this initial screen, we identified activities for twelve sulfatases (**Fig. 1c, Extended Data Figs. 3, 4** and **Supplementary Table 3**). Among these, 5 of the enzymes represent the first activities reported for their respective subfamilies: two S1_20 members (BT1636 and BT1622) were determined to target 3S-Gal, with BT1622 also preferentially cleaving 3S-*N*-acetyl-D-galactosamine (3S-GalNAc), two S1_16 enzymes (BT3796 and BT3057) cleave 4S-Gal/4S-GalNAc and one S1_46 enzyme (BT1918) cleaves 3S-GlcNAc, using the *N*-acetyl group as an absolute specificity determinant. This represents the first report of a bacterial sulfatase active on 3S-GalNAc, indicating that this sulfation could exist as a yet unidentified modification of host glycans. Subsequently, we refer to these enzymes by their gene/locus tag number with the corresponding activity in superscript (*e.g*., BT1636^3S-Gal^).

In addition to assigning new catalytic activities associated with three subfamilies that previously lacked any characterization, we also identified sulfatases displaying novel activities inside previously characterized subfamilies. These include three S1_15 enzymes (BT1624^6S-Gal/GalNAc^, BT3109^6S-Gal/GalNAc^ and BT4631^6S-Gal/GalNAc^) that extend this family, previously only known to include 6S-GalNAc sulfatases, to those cleaving 6S-Gal. Two members of S1_4 were active on 3S-Gal (BT4683^3S-Gal^) or 6S-Gal (BT3487^6S-Gal^), representing novel activities within this arylsulfatase subfamily. Finally, consistent with the activity previously described for S1_11 members, two enzymes were 6S-GlcNAc sulfatases (BT1628^6S-GlcNAc^ and BT3177^6S-GlcNAc^) (**Fig. 1c** and **Extended Data Fig. 3**).

Characterized sulfatases within the same subfamily, with the exception of two S1_4 members, cleaved the same sulfate ester linkages (**Fig. 1c**). However, despite these enzymes targeting the same linkages, their optimal activity depends on the surrounding glycan context. The activity of the 3S-Gal sulfatases is dependent on the linkage between Gal and GlcNAc with BT4683^3S-Gal^ showing a preference for 3’-sulfate-*N*-acetyl-D-lactosamine (3’S-LacNAc, 3’S-D-Gal-*β*1,4-D-GlcNAc). In contrast, BT1622^3S- Gal/GalNAc^ demonstrated enhanced activity towards 3’-sulfate-lacto-*N*-biose (3’S-LNB, 3’S-D-Gal-*β*1,3-D-GlcNAc) (**Extended Data Fig. 4a** and **Supplementary Table 3**).

Furthermore, additional affinity/activity studies revealed that BT1622^3S-Gal/GalNAc^ preferentially targets GalNAc and not Gal (**Extended Data Fig. 4b,c** and **Supplementary Table 3**), suggesting that this sulfatase evolved to optimally target sulfate *O*3-linked to GalNAc. *Bt* sulfatase activity is also affected by the presence of terminal epitopes such as those that occur in Lewis antigens. Despite BT1636^3S-Gal^ being equally active on 3’S-LacNAc and 3’S-LNB, this protein has a lower affinity and it is 100-fold less active when L-fucose (Fuc) is linked to GlcNAc (3’S-Lewis-a/x) (**Extended Data Fig. 4a,b** and **Supplementary Table 3**). While BT1622^3S-Gal/GalNAc^ is only weakly active on 3’S-Lewis-a antigen and not active at all on 3’S-Lewis-x, the reciprocal is true for BT4683^3S-Gal^ (**Supplementary Table 3**). The subfamily S1_15 enzyme BT1624^6S-Gal/GalNAc^ was only weakly active on 6S-Lewis-a/x antigens (**Extended Data Fig. 4a**), suggesting that these enzymes cannot accommodate Fuc linked to GlcNAc and that Fuc needs to be removed prior to sulfate cleavage. Additionally, affinity studies showed that BT3109^6S-Gal/GalNAc^ has a strong affinity for Gal, while the previously characterized GAG sulfatase BT3333^6S-GalNAc^ showed a preference for GalNAc, suggesting that optimal activity of S1_15 sulfatases likely depends on the glycan context and BT3333^6S-GalNAc^ evolved to target sulfated linkages in GAGs, a substrate that contains GalNAc but not Gal.

### Bt sulfatase activity on colonic mucin oligosaccharides (cMO)

We next tested the activity of *Bt* sulfatases on custom purified cMO, which we determined to contain at least 131 different oligosaccharides (**Supplementary Table 4)**. Only 4 of the 6 sulfatases tested displayed activity on cMOs (**Fig. 2a,b** and **Supplementary Table 4)**. BT1628^6S-GlcNAc^ and BT3177^6S-GlcNAc^ removed 6-*O*-sulfate from all GlcNAc structures but only when present at the non-reducing end of *O*-glycans confirming an exo-mode of action observed using commercial oligosaccharides (**Fig. 2** and **Extended Data Fig. 5a**). After incubation with BT1636^3S-Gal^, we were able to detect 14 new oligosaccharides and an overall increase of non-sulfated glycans (**Fig. 2** and **Extended Data Fig. 5b)**. Compared to the non-enzyme treated control, 36 oligosaccharides could no longer be detected after incubation with BT1636^3S-Gal^ (**Fig. 2a** and **Supplementary Table 4).** We determined the structures of 8 of these glycans and all present a terminal 3S-Gal (**Fig. 2)**. This sulfatase was active on 3’S-Gal-*β*1,3-GalNAc (core 1) and more complex sulfated structures built around other common core structures (**Fig. 2c)**, indicating that BT1636^3S-Gal^ evolved to accommodate the various linkages and substitutions found in mucin *O*-glycans. After incubation with BT4683^3S-Gal^, 9 oligosaccharides disappeared, indicating that this enzyme is active on a smaller subset of structures (**Fig. 2a** and **Supplementary Table 4)**. We were only able to determine the structure of one of those glycans and, unexpectedly, we found that this enzyme is endo-active on sialylated 3S-Gal (**Fig. 2b)**, which is consistent with the protein structural data discussed below. Additionally, the strong activity of BT1636^3S-Gal^ on larger glycans with unknown structures (**Supplementary Table 4)** reveals that porcine cMOs are highly sulfated at the O3-position of galactose. Although previous studies have reported the presence of 3S-Gal in colonic mucins^26–28^, the analysis of such complex samples is technically challenging and, until now, no precise enzymatic tools were available to probe these linkages. Together, these results represent the first report of HGM sulfatases active on colonic mucin *O*-glycans and highlight the possibility to use gut bacterial sulfatases as analytical tools in structural characterization of mucin glycans (see **Supplementary Discussion 2** for additional details of sulfatase activity on cMO).

**Figure 2.**
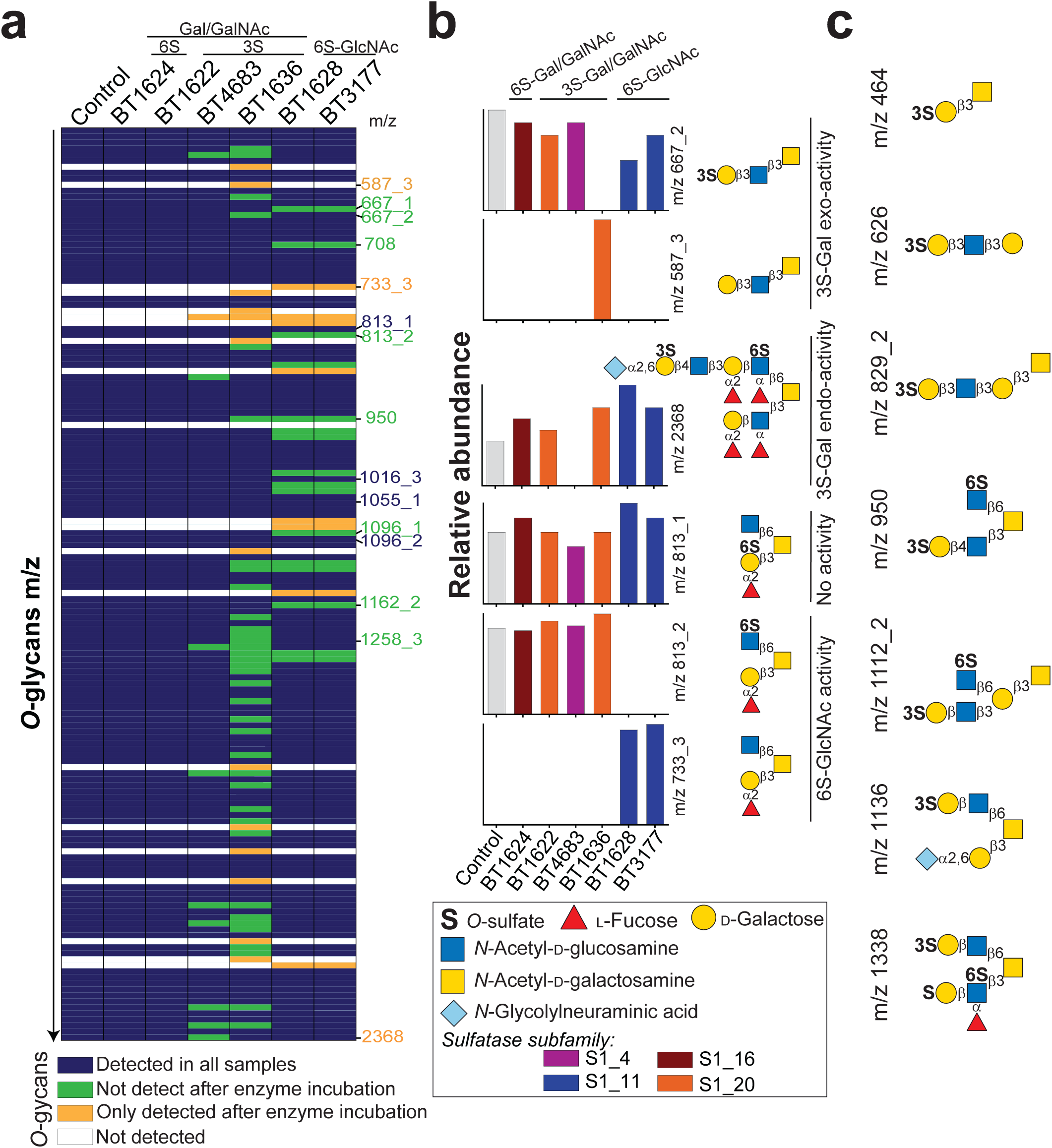
Activity of *Bt* sulfatases on colonic mucin *O*-glycans. **a,** Representation of *O*-glycans detected by mass spectrometry in cMO (control) and after sulfatase treatment from the lower (top) to the higher (bottom) mass range. **b,** Relative abundance and putative structures for the specific m/z shown in panel a. Remaining structures are shown in **Extended** figure 5a. **c,** Schematic representation of the putative structures that were not detected after treatment with BT1636^3S-Gal^.

### Structural characterization of 3S-Gal/GalNAc sulfatases

To further understand the molecular details of carbohydrate recognition by S1 sulfatases, we determined the crystal structures of the three different 3S-Gal sulfatases that belong to two subfamilies. Consistent with previous structures of S1 sulfatases, all 3 enzymes display a *α*/*β/a* topology with a C-terminal sub-domain, and the active site residues interacting with the sulfate group are fully conserved (**Extended Data Fig. 6a,b**). The structure of BT1636^3S-Gal^ in complex with the product LacNAc revealed that His177 coordinates with *O*4 of Gal and mutation to alanine ablates enzyme activity, suggesting that this residue is the major specificity determinant for Gal (presenting an axial *O*4) over glucose (equatorial *O*4) (**Fig. 3** and **Supplementary Table 3**). BT1636^3S-Gal^ also makes strong interactions with O2 via R353 and E334. The essential His177 in BT1636^3S-Gal^ is highly conserved (92%) within S1_20 sulfatases (**Extended Data Fig. 7**). In BT1622^3S- Gal/GalNAc^, mutation of the corresponding H176 to alanine causes a ∼300-fold reduction in activity (**Supplementary Table 3**), further highlighting the importance of this residue in Gal recognition. In BT1622^3S-Gal/GalNAc^, R353 and E334 are replaced by C357 and N334, amino acids that present shorter side chains that allow the accommodation of a *C*2-linked *N*-acetyl group found in GalNAc (**Fig. 3**). This observation is consistent with the ability of BT1622^3S-Gal/GalNAc^ to cleave 3S-GalNAc and preferentially bind GalNAc over Gal, while BT1636^3S-Gal^ only recognizes Gal (**Extended Data Fig. 4b,c**). Additionally, and consistent with *exo*-activity observed for both enzymes, the substrates are buried in a deep pocket and only *O1* is solvent exposed (**Fig. 3**).

**Figure 3.**
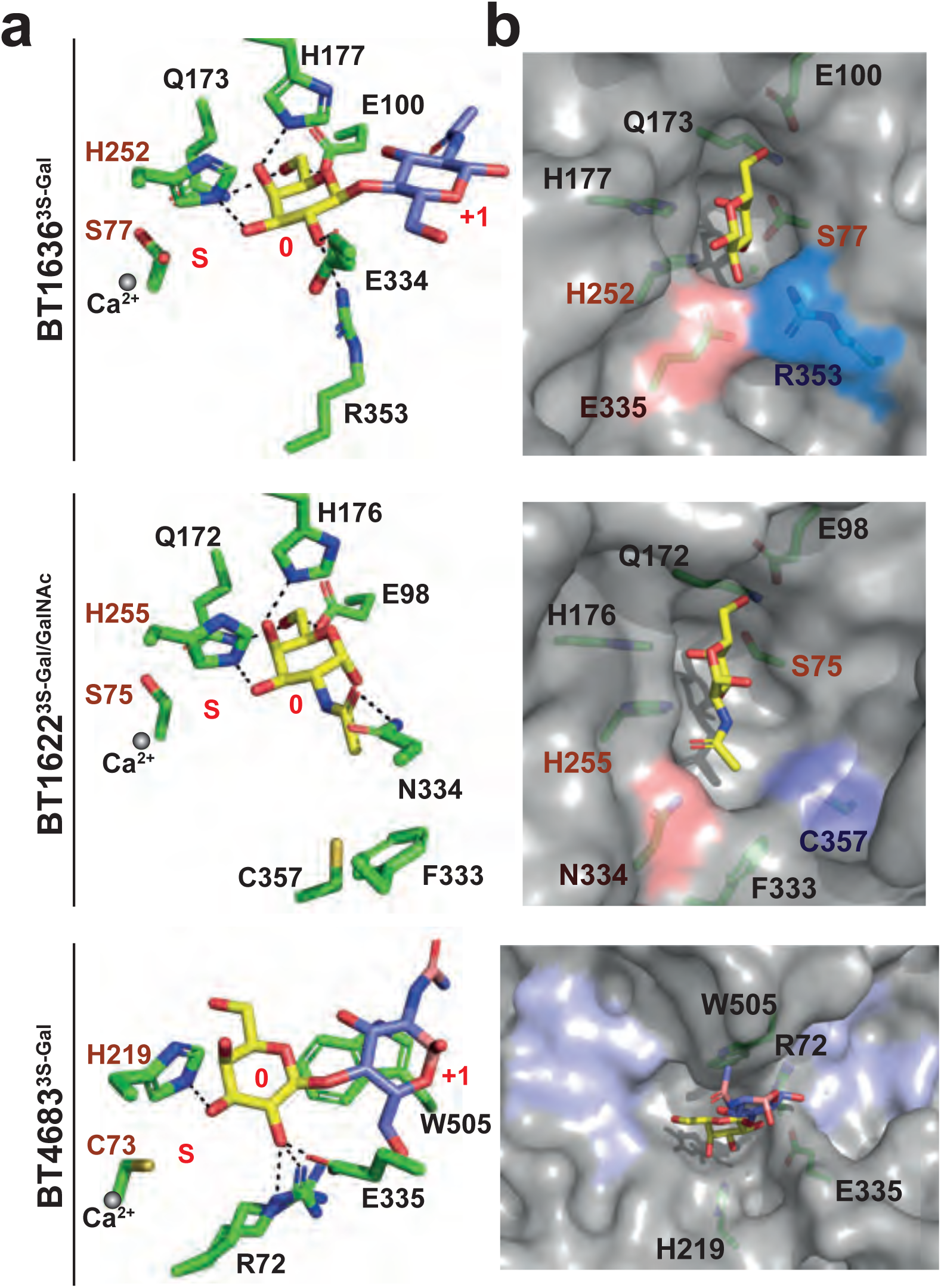
Crystal structures of 3S-Gal/GalNAc sulfatases. **a,** Schematic representation of the residues interacting with targeted sugars, including the putative catalytic residues (in dark red), the calcium ion (grey sphere) and subsites S, 0 and +1 highlighted in red. BT1636^3S-Gal^ and BT4683^3S-Gal^ in complex with LacNAc (D-Gal-*β*1,4-D-GlcNAc) and BT1622^3S-Gal/GalNAc^ in complex with GalNAc. **b,** Surface representation of the active pocket. The equivalent Gal/GalNAc specificity residues in BT1636^3S-Gal^ and BT4683^3S-Gal^ are highlighted in red and blue. The open active site of BT4683^3S-Gal^ is highlighted in purple. In all structures the amino acids and ligands are represented as stick.

For the S1_4 enzyme BT4683^3S-Gal^, a structure solved in complex with LacNAc did not reveal any interaction between the protein and the O4 of Gal. In BT4683^3S-Gal^, the interaction with Gal is driven by the residues R72 and E335, spatially equivalent to R353 and E334 in BT1636^3S-Gal^, that form hydrogen bonds with *O2* of D-Gal (**Fig. 3**) and disruption of either of these residues eliminates activity (**Supplementary Table 3**). However, in BT4683^3S-Gal^, a sulfatase that does not have any affinity for monosaccharides (**Extended Data Fig. 4b**), the active site is located in an open cleft (**Fig. 3**) that allows the accommodation of additional substitutions on Gal (**Extended Data Fig. 6d**). This finding is consistent with the apparent endo-activity found using cMO. Together, these structures reveal the key specificity determinants in 3S-Gal/GalNAc sulfatases, highlighting that these enzymes have evolved to target sulfate groups in the different contexts in which they are found in complex host glycans. This is especially true for BT1636^3S-Gal^ which utilizes high affinity interactions with both O2 and O4 to drive enhanced activity to remove terminal 3S-Gal linkages in cMOs (see **Supplementary Discussion 3** for additional details of sulfatase structural characterization and phylogeny).

### Roles of sulfatases in B. thetaiotaomicron O-glycan utilization

*Bt* is able to utilize cMO as a sole carbon source (**Fig. 1b**), but the key enzymes involved in the degradation of these glycans remain unclear. Highlighting the importance of sulfatases to this symbiont’s physiology, deletion of the gene encoding the only anaerobic sulfatase maturating enzyme (*anSME*) eliminates activation of all 28 S1 sulfatases^15^ and the ability of *Bt* to grow efficiently on cMOs (**Fig. 4a**). Based on this observation we generated a series of strains with compounded gene deletions in which one or several groups of sulfatases were eliminated based on their biochemical activity. Deletion of all 3S-Gal/GalNAc sulfatases (*Δbt1636^3S-Gal^ + Δbt1622^3S-Gal/GalNAc^* + *Δbt4683^3S-Gal^*) resulted in a growth phenotype similar to *ΔanSME* (**Extended Data Fig. 8a**). Interestingly, we observed a similar growth defect when just BT1636^3S-Gal^ was deleted, but not the other 3S-Gal sulfatases (**Fig. 4a** and **Extended Data Fig. 8a**), consistent with the prominent activity of the recombinant form of this enzyme on cMOs. In contrast, a strain with compounded deletions of eight other sulfatases (*Δbt1622^3S-Gal/GalNAc^* + *Δbt4683^3S-Gal^* + *Δbt1624^6S-Gal/GalNAc^* + *Δbt3109^6S-Gal^* + *Δbt4631^6S-Gal/GalNAc^* + *Δbt1628^6S-GlcNAc^* + *Δbt3177^6S-GlcNAc^* + *Δbt3051^putative_6S-GlcNAc^*) displayed a growth phenotype similar to wild-type (**Fig. 4a**), indicating that these enzymes are not essential for cMO utilization. However, a *Δ10X sulf* mutant, which included the deletion of BT1636^3S-Gal^ and the two 4S-Gal/GalNAc sulfatases, showed a similar growth defect as *ΔanSME* and *Δbt1636*. Complementation of this and other loss of function mutants with only *bt1636^3S-Gal^* restored growth on cMO to levels similar to wild-type (**Fig. 4a** and **Extended Data Fig. 8a**). Cellular localization experiments revealed that BT1636*^3S-Gal^* is located at the cell surface in *Bt* (**Fig. 4b**) and together these data suggest that this single 3S-Gal sulfatase is a critical cell surface enzyme involved in the utilization of sulfated *O*-glycans that are prominent in the colon.

**Figure 4.**
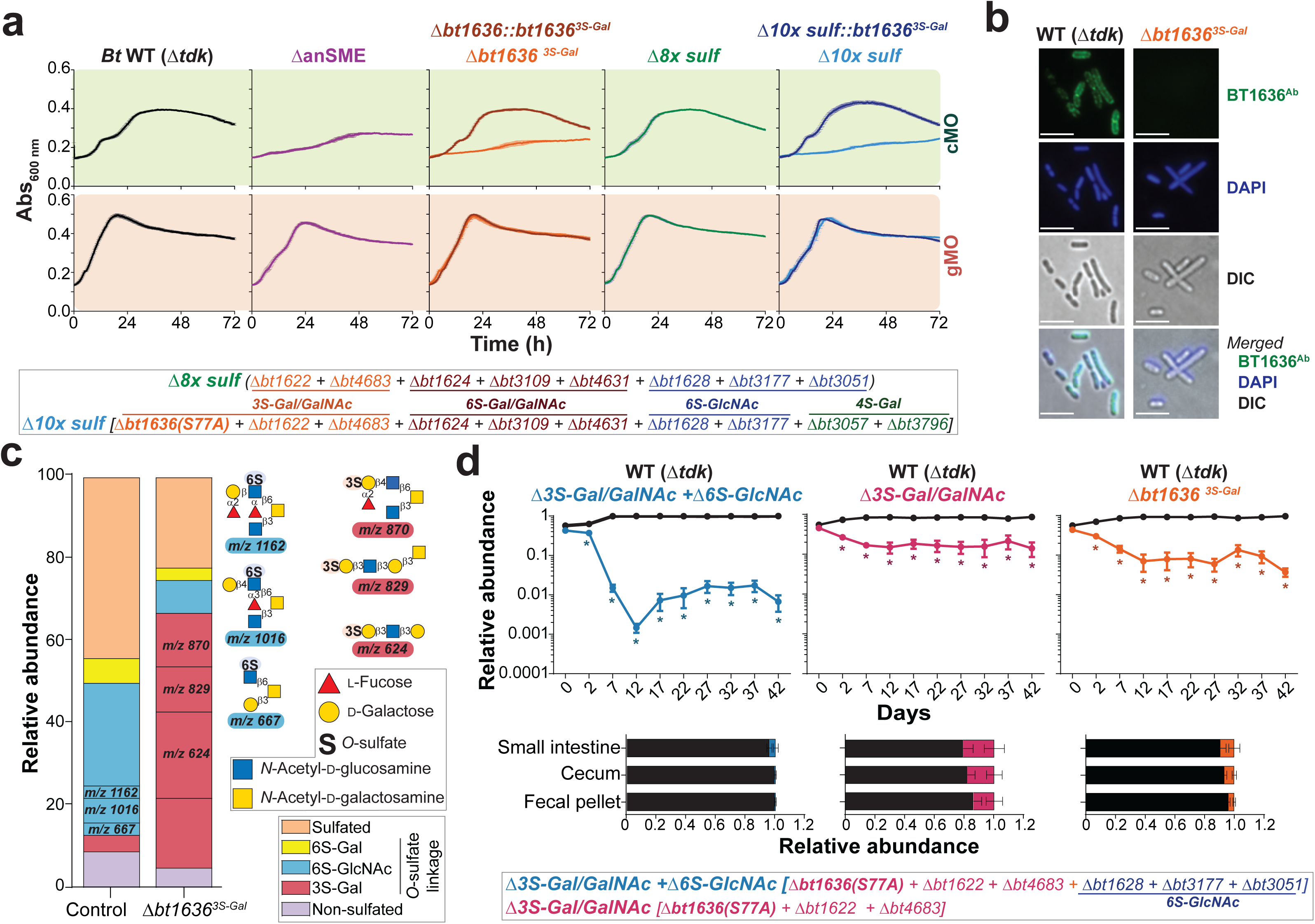
BT1636^3S-Gal^ activity is required for the utilization of cMO and competitive fitness *in vivo*. **a**, Growth of *Bt* wild-type *Δtdk* (WT), different sulfatase gene-deletion mutants (named “*ΔbtXXXX”*) and strains complemented with *bt1636^3S-Gal^* on colonic or gastric mucin O-glycans (cMO and gMO, respectively) (line represents the average of biological replicates (n = 3) and error bars denote s.e.m.) **b,** Immunofluorescent and differential interference contrast (DIC) microscopy of *Bt* WT and sulfatase mutant staining with polyclonal antibody (Ab) against BT1636^3S-Gal^ (green) and DNA staining with DAPI (blue). **c,** Relative abundance of different *O*-glycans detected by mass spectrometry in *Δbt1636^3S-Gal^* culture supernatant or cMO in minimal media without bacteria (control), after 96h in anaerobic conditions. The mass and associated structure of the 3 more abundant glycans in both samples are shown. **d,** *in vivo* competitions in gnotobiotic mice (n = 5-9 separate mice from two separate experiments, except competition of the *Δ6S-GlcNAc* mutant that showed no defect in one experiment) fed fiber-free diet and inoculated with WT and mutants. The fecal relative abundance of each strain was determined along the time course and in small intestine and cecum at day 42 (experimental endpoint). The relative abundance in each mouse is represented in the respective light colour. The error bars denote s.e.m. Significant differences between wild-type and mutant strain were compared at each time point using student’s t-test (paired, one tail) and * indicates sample days in which the mutant was significantly different (p < 0.01) from the wild-type.

To further investigate the role of BT1636^3S-Gal^ in *O*-glycan utilization, we analyzed the oligosaccharides present in the culture supernatant of the wild-type and *Δbt1636^3S-Gal^* strains after growth on cMO. Consistent with a robust ability of *Bt* to degrade diverse colonic *O*-glycans, no oligosaccharides were detected in wild-type supernatant (**Supplementary Table 5**). Compared to the cMO used as substrate (control), the supernatant of *Δbt1636^3S-Gal^* showed a 20-fold accumulation of terminal 3S-Gal capped glycans, suggesting that these could not be degraded. Indeed, the three most common 3S-Gal structures detected in the *Δbt1636^3S-Gal^* supernatant accounted for 50% of the total oligosaccharides detected in this sample (**Fig. 4c**). These results reveal that deletion of BT1636^3S-Gal^ results in loss of the ability to utilize 3S-Gal *O*-glycans. Interestingly, 49 of the 72 glycans detected in *Δbt1636^3S-Gal^* supernatant were not present in the control sample (**Supplementary Table 5**), suggesting that although the mutant does not grow well on cMO it is able to modify these oligosaccharides to generate new glycans (**Supplementary discussion 5**). These data, combined with the cell surface location of BT1636^3S-Gal^, suggest that this sulfatase is required early in *O*-glycan catabolism likely by cleaving 3S-Gal from *O*-glycans prior to importing them into the periplasm where these oligosaccharides will be sequentially degraded by additional enzymes and also serve as cues for activating transcription of the *O*-glycan PULs (**Extended Data Fig. 2)**. Although *Bt* does encode two additional 3S-Gal/GalNAc sulfatases, the low activity of these additional sulfatases on cMOs (**Fig. 2**) and their likely periplasmic location, suggests why these enzymes cannot compensate for loss of BT1636^3S-Gal^. Interestingly, all of the *Bacteroides* species able to utilize cMO (**Fig. 1a**) have homologues of BT1636^3S-Gal^, suggesting that this activity plays a key role in mucin utilization by other HGM members (**Supplementary Table 6**).

Finally, to investigate the requirement for specific sulfatases *in vivo*, we utilized gnotobiotic mice in which we competed individual mutants against the wild-type strain to evaluate their colonization fitness. It has been reported that mouse colonic Muc2 prominently displays 6S-GlcNAc modifications^29^. However, mutants lacking either just the two active 6S-GlcNAc sulfatases (*Δ6S-GlcNAc* double mutant), or these two enzymes, another putative 6S-GlcNAc sulfatase and all three 6S-Gal/GalNAc sulfatases (*Δ6S-GlcNAc* + *Δ6S-Gal/GalNAc* hexa mutant), competed equally with wild-type (**Extended Data Fig. 8c**), suggesting that neither of these two sulfatase activities are essential determinants *in vivo*. A significant defect was observed with a mutant lacking all 3S-Gal/GalNAc sulfatases (**Fig. 4d**). The fitness defect was exacerbated by eliminating 3S-Gal/GalNAc and 6S-GlcNAc sulfatase activities together (*Δ3S-Gal/GalNAc* + *Δ6S-GlcNAc*) (**Fig. 4d**), suggesting that they synergize *in vivo*. Consistent with its prominent role in cMO utilization *in vitro*, a mutant lacking just BT1636^Gal-3Sulf^ displayed a significant defect that was similar or slightly more severe compared to the *Δ3S-Gal/GalNAc* mutant when competed with the wild-type strain (**Fig. 4d**), further suggesting that this enzyme plays an essential role in gut colonization by allowing *Bt* to access 3S-Gal *O*-glycans (see **Supplementary Discussion 4-5** for additional details regarding growth and competition of mutants *in vitro* and *in vivo*).

## Conclusion

To degrade the complex *O*-glycans found in mucins some HGM bacteria have evolved complex arsenals of degradative enzymes which include diverse sulfatases. Disarming all of the sulfatases in *Bt* via *anSME* deletion results in drastically reduced competitive colonization (**Extended Data Fig. 8c**)^15^ and an inability to elicit colitis in an animal model of IBD^16^. While these findings support a critical role for active sulfatases in both fitness and promoting inflammation, they provide no insight into the complexity of catalytic modifications carried out by these enzymes. In this study, we reveal that *Bt* has a robust ability to grow on highly sulfated mucin oligosaccharides from colonic tissue and that it possesses active sulfatases capable of removing sulfate groups in all contexts in which sulfation is known to occur in mucin, including novel specificities. Surprisingly, we found that a single key sulfatase is essential for growth on colonic mucin *O*-glycans. This cell surface enzyme removes 3-sulfate capping Gal allowing the degradation of these glycans by additional enzymes. This critical role of BT1636^3S-Gal^ supports the conclusion that keystone steps exist in the complex pathway of mucin degradation. Further delineation of these critical steps, along with identification of the corresponding enzymes(s), are a prerequisite to modulating such events and potentially inhibiting mucin-degrading activities in bacteria that contribute to disease.

## Methods

### Recombinant Protein Production

Genes were amplified by PCR using the appropriate primers and the amplified DNA cloned in pET28b using *NheI/XhoI* restriction sites or pETite (Expresso^TM^ T7 cloning and expression system, Lucigen) generating constructs with either N- or C-terminal His_6_ tags (**Supplementary Table 7**). The catalytic serine was mutated to cysteine since *Escherichia coli* in only able to convert cysteine to formylglycine. Recombinant genes were expressed in *Escherichia coli* strains BL21 (DE3) or TUNER (Novagen), containing the appropriate recombinant plasmid, and cultured to mid-exponential phase before induction with 1 mM (BL21(DE3)) or 0.2 mM (TUNER) of isopropyl β-D-1-thiogalactopyranoside; cells were cultured for another 16 h at 16°C and 180 rpm. Recombinant protein were purified to >90% electrophoretic purity by immobilized metal ion affinity chromatography using a cobalt-based matrix (Talon, Clontech) and eluted with imidazole as described previously^24^. For the proteins selected for structural studies, another step of size exclusion chromatography was performed using a Superdex 16/60 S200 column (GE Healthcare), with 10 mM HEPES, pH 7.5, and 150 mM NaCl as the eluent, and they were judged to be ≥95% pure by SDS-PAGE. Protein concentrations were determined by measuring absorbance at 280 nm using the respective molar extinction coefficient. When necessary, proteins were then concentrated by centrifugaton using a molecular mass cutoff of 30 kDa.

### Site-Directed Mutagenesis

Site-directed mutagenesis was conducted using the PCR-based QuikChange kit (Stratagene) and conducted according to the manufacturer’s instructions, using the appropriate plasmid as the template and primers (**Supplementary Table 8**). All mutations were confirmed by DNA sequencing.

### Sources of purified carbohydrates

All carbohydrates were from Sigma, Carbosynth or Dextra Laboratories. All other chemical reagents were purchased from Sigma. The 3S-GalNAc was chemically synthesized as previously described^30^.

### Mucin purification

Gastric mucin oligosaccharides (gMO) were purified from commercial available porcine gastric mucins (type III, Sigma) as previously described^14^. Colonic mucins oligosaccharides (cMO) were purified from pig distal pig colons and rectum. Briefly, the tissue was open and the fecal contents were carefully removed. The mucosa was scrapped off and mucus was extracted by homogenizing the tissue in at least 5 times volume of extraction buffer (6 M guanidine chloride, 5 mM EDTA, 10 mM NaH_2_PO_4_, pH 6.5) and slow stirring at 4°C for 16 h. The solution was spun down at 15,000 rpm and 10°C for 30 min and supernatant was discharged. The pellets were resuspended in extraction buffer and the process was repeated until the supernatant was clear for at least two extractions. After the extraction the mucins were solubilized by reducing the disulfide bonds. The pellets were resuspended in fresh reduction buffer (6 M guanidine chloride, 0.1 M Tris, 5 mM EDTA, pH 8.0) containing 25 mM of 1,4-dithiothreitol and slowly stirred at 37°C for 5 h. After this incubation, 62.5 mM of iodoacetamide were added and the solution was stirred slowly in the dark at room temperature for 16 h. The solution was centrifuged at 10,000 rpm at 4°C for 30 min and the supernatant containing the solubilized mucins was extensively dialysed into water. Samples were dissolved into 100 mM Tris-HCl pH 8.0 containing 1 mg/ml of trypsin and incubated slowly stirring at 37°C for 16 h. The glycans were *β*-eliminated by adding 0.1 M NaOH and 1 M NaBH_4_ and incubate the solution at 65°C for 18 h. After cooling the solution to room temperature, the pH was adjusted to 7.0 with concentrated HCl and extensively dialysed in water. The released porcine colonic mucin glycans were recovered by lyophilization the solution until completely dry and used in further experiments.

### HPLC and TLC sulfatase enzymatic assays

The sulfatase activity screen against commercially available sulfated oligosaccharides (**Supplementary Table 2**) was performed with 1 *μ*M of recombinant enzyme and 1 mM of substrate in 10 mM MES pH6.5 with 5 mM CaCl2 for 16h at 37°C. Sulfated *N*-acetyl-D-lactosamine and lacto-*N*-biose were generated by incubating the respective sulfated Lewis antigens with 1 μM of *α*-1,3/1,4-fucosidase BT1625^31^ in the same conditions. Reactions were analysed by thin layer chromatography (TLC). Briefly, 2 μL of each sample was spotted onto silica plates and resolved in butanol:acetic acid:water (2:1:1) running buffer. The TLC plates were dried, and the sugars were visualized using diphenylamine stain (1 ml of 37.5% HCl, 2 ml of aniline, 10 ml of 85% H3PO3, 100 ml of ethyl acetate and 2 g diphenylamine) and heated at 100°C for 20 min. When relevant, the enzymatic activity was confirmed by high-performance anionic exchange chromatography (HPAEC) with pulsed amperometric detection using standard methodology. The sugars (reaction substrate/products) were bound to a Dionex CarboPac P100 column and eluted with an initial isocratic flow of 10 mM NaOH during 20 min then a gradient of 10-100 mM of NaOH for 20 min at a flow rate of 1.0 ml min^-1^. The reaction products were identified using the appropriated standards. All experiments were performed in triplicate.

### Liquid Chromatograph-Electrospray Ionization Tandem Mass Spectrometry

Enzymatic reactions of sulfatases in colonic mucin oligosaccharides and culture supernatant were cleaned up with graphitized carbon^32^. Reactions with sulfated defined saccharides were reduced and desalted. Briefly, reactions were dried in Speed vac, reconstituted in 20 *μ*L of 50 mM NaOH and 500 mM NaBH_4_ and incubated at 50°C for 3 h. Reactions were cool down on ice, neutralized with 1 *μ*L of glacial acetic acid and desalted using a cation exchange column containing AG^®^50W-X8 resin. All cleaned and desalted reactions were reconstituted in water before analysis by liquid chromatograph-electrospray ionization tandem mass spectrometry (LC-ESI/MS). The oligosaccharides were separated on a column (10 cm × 250 *μ*m) packed in-house with 5 µm porous graphite particles (Hypercarb, Thermo-Hypersil, Runcorn, UK). The oligosaccharides were injected on to the column and eluted with a 0-40 % acetonitrile gradient in 10 mM ammonium bicarbonate over 46 min at a flow rate of 10 *μ*l/min.. A 40 cm × 50 µm i.d. fused silica capillary was used as transfer line to the ion source. Samples were analyzed in negative ion mode on a LTQ linear ion trap mass spectrometer (Thermo Electron, San José, CA), with an IonMax standard ESI source equipped with a stainless steel needle kept at –3.5 kV. Compressed air was used as nebulizer gas. The heated capillary was kept at 300°C, and the capillary voltage was –33 kV. Full scan (*m/z* 380-2,000, two microscan, maximum 100 ms, target value of 30,000) was performed, followed by data-dependent MS^2^ scans (two microscans, maximum 100 ms, target value of 10,000) with normalized collision energy of 35%, isolation window of 2.5 units, activation q*=*0.25 and activation time 30 ms). The threshold for MS^2^ was set to 300 counts. Data acquisition and processing were conducted with Xcalibur software (Version 2.0.7). Glycans were identified from their MS/MS spectra by manual annotation and validated by available structures stored in Unicarb-DB database (2020-01 version) (**Supplementary Fig. 1**)^33^. *O*-Glycan structural characterization was based on diagnostic fragment ions^34^.The schematic glycosidic or cross-ring cleavages were assigned according to the Domon and Costello nomenclacture^35^. For comparison of glycan abundance between samples, the individual glycan structures were quantified relative to the total content by integration of the extracted ion chromatogram peak area using Progenesis QI. The area under the curve (AUC) of each structure was normalized to the total AUC and expressed as a percentage.

### Microfuidic-based enzymatic desulfation assays

Sulfated carbohydrates were labelled at their reducing end with BODIPY which has a maximal emission absorbance of ∼503 nm, which can be detected by the EZ Reader via LED-induced fluorescence. Non-radioactive mobility shift carbohydrate sulfation assays were optimised in solution with a 12-sipper chip coated with CR8 reagent using a PerkinElmer EZ Reader II system using EDTA-based separation buffer. This approach allows real-time kinetic evaluation of substrate de-sulfation^36^. Pressure and voltage settings were adjusted manually (1.8 psi, upstream voltage:2250 V, downstream voltage: 500 V) to afford optimal separation of the sulfated product and unsulfated substrate with a sample (sip) time of 0.2 s, and total assay times appropriate for the experiment. Individual de-sulfation assays were carried out at 28°C after assembly in a 384-well plate in a final volume of 80 μl in the presence of substrate concentrations between 0.5 and 20 *μ*M with 100 mM Bis-Tris-Propane, 150 mM NaCl, 0.02% (v/v) Brij-35 and 5 mM CaCl_2_. The degree of de-sulfation was calculated by peak integration using EZ Reader software, which measures the sulfated carbohydrate : unsulfated carbohydrate ratio at each individua time-point. The activity of sulfatase enzymes was quantified in ‘kinetic mode’ by monitoring the amount of unsulfated glycan generated over the assay time, relative to control assay with no enzyme; with sulfate loss limited to ∼20% to prevent of substrate and to ensure assay linearity. *k*_cat_/*K*_M_ values, using the equation V_0_=(V_max_/*K*_M_)/S, were determined by linear regression analysis with GraphPad Prism software. Substrate concentrations were varied to ensure assay linearity, and substrate concentrations present were significantly <*K*_M_.

### NMR desulfation assays

NMR experiments, monitoring the de-sulfation of 6S-D-galactose and 6S-*N*-acetyl-D-galactosamine, were conducted in D_2_O with 50 mM sodium phosphate, pH 7.0, supplemented with 150 mM NaCl at 25°C on a 800MHz Bruker Avance III spectrometer equipped with a TCI CryoProbe and a 600MHz Bruker Avance II+ spectrometer, also fitted with a TCI CryoProbe. 1D and 2D proton and TOCSY spectra (mixing time 80 ms) were measured using standard pulse sequences provided by the manufacturer. Spectra were processed and analysed using TopSpin 3.4A and TopSpin 4.0 software (Bruker). Galactose integrals were recorded directly for the C(6)H_2_-OH peak within the region 3.694 to 3.721ppm, referenced to the combined C(2) peaks of D-galactose and 6S-D-galactose with in the region 3.415 to 3.475ppm. Similarly, 6S-*N*-acetyl-D-galactosamine integrals were recorded directly for the C(6)H_2_-OH peak within the region 3.674 to 3.747ppm, referenced to the combined C(4) peaks for *N*-acetyl-D-galactosamine and 6S-*N*-acetyl-D-galactosamine in the region 3.925 to 3.968ppm.

### Differential scanning fluorimetry

Thermal shift/stability assays (TSAs) were performed using a StepOnePlus Real-Time PCR machine (LifeTechnologies) and SYPRO-Orange dye (emission maximum 570 nm, Invitrogen) as previously described^37^ with thermal ramping between 20 and 95°C in 0.3°C step intervals per data point to induce denaturation in the presence or absence of various carbohydrates as appropriate to the sulfatase being analysed. The melting temperature (Tm) corresponding to the midpoint for the protein unfolding transition was calculated by fitting the sigmoidal melt curve to the Boltzmann equation using GraphPad Prism, with R^2^ values of >0.99. Data points after the fluorescence intensity maximum were excluded from the fitting. Changes in the unfolding transition temperature compared with the control curve (ΔT_m_) were calculated for each ligand. A positive ΔT_m_ value indicates that the ligand stabilises the protein from thermal denaturation, and confirms binding to the protein. All TSA experiments were conducted using a final protein concentration of 5μM in 100 mM Bis-Tris-Propane (BTP), pH 7.0, and 150 mM NaCl supplemented with the appropriate ligand. When BT1622^3S-Gal/GalNAc^ and BT1636^3S-Gal^ were assessed against 3’S-LacNAc and 3’S-LNB 100 mM Hepes (pH 7.0) was employed instead of BTP, although no difference in the T_m_ value of the proteins was observed. Three independent assays were performed for each protein and protein ligand combination.

### Glycan labelling

Sulfated saccharide samples were labelled according to a modification of the method previously described reporting the formation of N-glycosyl amines for 4,6-O-benzilidene protected D-gluopyranose monosaccharides with aromatic amines^38^. Briefly, the lyophilised sugar (1 mg) was dissolved in 0.50 ml anhydrous methanol in a 1.5 ml screw-top PTFE microcentrifuge tube. 0.1 mg, BODIPY-FL hydrazide (4,4–difluoro-5,7– dimethyl-4-bora-3a,4a–diaza-s-indacene-3-propionic acid hydrazide, *λ*_ex./em. 505/513_, extinction coefficient 80,000 M^-1^ cm^-1^) was added and the mixture vortexed (1 min), then incubated in darkness at 65°C for 24 h. The products were then cooled and a portion purified by TLC on silica coated aluminium plates and developed with methanol or 1:1 v/v ethyl acetate/methanol to provide R_f_ values suitable to allow separation of unreacted label from labelled glycan product. The unreacted BODIPY-FL label (orange on the TLC plate) was identified by reference to a lane containing the starting material (BODIPY-FL hydrazide) on the TLC plate, allowing differentiation from the putative labelled product (also orange). This latter band was scraped from the plates and extracted in fresh methanol (2 x 0.5 ml), spun for 3 min at 13,000 x *g* and the supernatant was recovered and dried (rotary evaporator) to recover the fluorescent-coloured product (bright green when dissolved in aqueous solution), which was then employed in subsequent experiments.

### Anaerobic bacterial culture and genetic manipulation

All strains were anaerobic grown at 37 °C in a chamber (10% H_2_, 5% CO_2_, and 85% N_2_; Coy Manufacturing, Grass Lake, MI). *Bacteroides* type strains were culture in either tryptone-yeast extract-glucose medium (TYG), brain heart infusion medium or minimal medium (MM) containing an appropriate carbon source. *Bacteroides massilliensis* and *Akkermansia muciniphila* were culture as described before^21, 39^. *Bt* strains containing specific gene deletions or inactivated versions of enzymes (BT1636^3S-Gal^ S77A) were made by counterselectable allelic exchange as previously described^40^. Complemention of deletion strains was performed using pNBU2 vector as previously described^40^, containing a constitutive promotor used previously^41^. All primers used to generate the mutants and complementation are listed in **Supplementary Table 9**. Growth of the WT and mutants was measured on an automated plate reader by increase in absorbance at 600 nm in 96-well plates containing 200 *μ*l of minimal media mixed with the respective filter-sterilised (monosaccharide and gMO) or autoclave-sterilised cMO as described before^39^. To achieve consistent growth, all carbon sources were used at 5 mg/ml with exception of gMO that was added in a final concentration of 10 mg/ml. All growth curves presented are averages and s.e.m of three technical replicates.

### Crystallization of carbohydrate sulfatases

After purification, all proteins were carried forward in the same eluent as used for the size exclusion chromatography (see Recombinant Protein Production). Sparse matrix screens were set up in 96-well sitting drop TTP Labtech plates (400-nL drops). Initial hits crystals for all proteins were obtained between 20 and 35 mg/ml. For BT1622^3S-Gal/GalNAc^ and BT1636^3S-Gal^ wildtype *Bt* variants were used, having a Ser at the catalytic formylglycine position, whilst for for BT4683^3S-Gal^ the S73C mutant was used. BT1622^3S-Gal/GalNAc^ with 20 mM LNB crystallised in 20% PEG 3350 and 0.2 M sodium citrate tribasic dihydrate. BT1636^3S-Gal^ with 20 mM LacNAc crystallised in 40% MPD and 0.2 M sodium cacodylate pH 6.5. for BT4683^3S-Gal^ with 20 mM LacNAc crystallised in 20% PEG 3350, 0.2 M sodium iodide and BTP pH 8.5. All crystals were cryoprotected with the addition of the ligand they were crystallised with plus 20% PEG 400 and 20% glycerol was used as the cryoprotectant for BT4683^3S-Gal^ and BT1622^3S-Gal/GalNAc^, respectively. No cryoprotectant was added to BT1636^3S-Gal^ crystals. Data were collected at Diamond Light Source (Oxford) on beamlines I03, I04, I04-1 and I24 at 100 K. The data were integrated with XDS^42^, or Xia2^43^ 3di or 3dii and scaled with Aimless^44^. Five percent of observations were randomly selected for the R_free_ set. The phase problem was solved by molecular replacement using the automated molecular replacement server Balbes^45^ for all proteins except BT1622^3S-Gal/GalNAc^. The phase problem for BT1622^3S-Gal/GalNAc^ was initially solved using Molrep^46^ and BT1636^3S-Gal^ as the search model. This gave a partial solution, which could not be fully solved due to twinning. An acceptable model of BT1622^3S-Gal/GalNAc^ was constructed to be used to better solve the phase problem and the molecular replacement was re-performed. Models underwent recursive cycles of model building in Coot^47^ and refinement cycles in Refmac5^48^. Bespoke ligands were generated using JLigand^49^. The models were validated using Coot^47^ and MolProbity^50^. Structural figures were made using Pymol and all other programs used were from the CCP4^51^ and CCP4i2 suite^52^. The data processing and refinement statistics are reported in **Supplementary Table 10**.

### Immunolabelling of BT1636 in *Bt* cell surface

For the fluorescence microscopy, *Bt* cells (Wild type (*Δtdk*) and *Δbt1636^3S-Gal^*) were grown to early exponential phase (Abs_600nm_ 0.25–0.35) in rich TYG medium. One ml of the cultures was collected, centrifuged at 13,000 x *g*, and subsequently washed three times in MM with no carbon source. *Bt* cells incubated with cMO for four hours and fixed in 4.5% formalin overnight at 4°C with gentle rocking. Cells were stained with a polyclonal antibody raised in rabbit against purified recombinant BT1636 (BT1636^Ab^, Cocalico Biologicals) and detected with an Alexa Fluor^®^ 488-conjugated goat anti-rabbit IgG secondary antibody (Molecular Probes). Images were taken with Zeiss Apotome using the same exposure time between samples.

### Gnotobiotic Mouse Experiments

All experiments involving animals, including euthanasia via carbon dioxide asphyxiation, were approved by the University Committee on Use and Care of Animals at the University of Michigan (NIH Office of Laboratory Animal Welfare number A3114-01) and overseen by a veterinarian. Groups of 3 to 5, 6-8 week old germfree Swiss Webster mice were randomly assigned to each experiment. 7 days prior gavage the animals diet was switched to a fiber-free diet (Envigo-Teklad TD 130343) that was maintained through all the experiment. At day 0, mice were gavage with equal amount of *Bt* WT strain and mutant and fecal samples were collected at day 2 and every 5 days until day 42. At the end-point of the experiment distal small intestine and cecal contents were also collected. The bacteria gDNA extraction and quantification by qPCR of the relative abundance of each strain on the various samples was carried out as described previously^39^.

### Phylogenetic analysis

To maximise sequence coverage, and avoid repetition, we selected 800 and 920 representative sequences of subfamily S1_20 (composed of 1356 sequences) and S1_4 (composed of 1895 sequences), respectively. The sequences were aligned by MAFFT v.7^53^ using L-INS-i algorithm. The multiple sequence alignment was visualized by Jalview software v.11.0^54^ and non-aligned regions were removed. 404 and 364 positions were used for the S1_4 and S1_20 phylogeny, respectively. Phylogeny was made using RAxML v. 8.2.4^55^. The phylogenetic tree was build with the Maximum Likelihood method^56^ and the LG matrix as evolutive model^57^ using a discrete Gamma distribution to model evolutionary rate differences among sites (4 categories). The rate variation model allowed for some sites to be evolutionarily invariable. The reliability of the trees was tested by bootstrap analysis using 1,000 resamplings of the dataset^58^. All the final global phylogenetic trees were obtained with MEGA v.7^59^. Fifteen S1_0 sequences from the sulfAtlas database were used as an outgroup.

### Quantification and statistical analysis

For *in vivo* competitions, when three or more fecal samples were collected, Student’s t tests (one-tailed, paired) were performed for each time point in Excel. When necessary, the statistical analysis for remaining samples in stated in the respective figure legend.

### Data availability statement

Source Data for all experiments, along with corresponding statistical test values, where appropriate, are provided within the paper and in Supplementary information. The crystal structure dataset generates have been deposit in the in the Protein Data Bank (PDB) under the following accession numbers: 7ANB, 7ANA, 7AN1 and 7ALL. The LC-MS/MS raw files and annotated structures are submitted to the Glycopost (https://glycopost.glycosmos.org/preview/12430260615f9d5733a1a5d, code 1955) and Unicarb-DB database, respectively.

### Code availability statement

No new codes were developed or compiled in this study

## Supporting information

Supplementary Table 4, provided separately, multiple tabs

Supplementary Table 5, provided separately, multiple tabs

## Acknowledgements

This project has received funding from the European Union’s Horizon 2020 research and innovation programme under the Marie Skłodowska-Curie grant agreement N° 748336. This work was supported by National Institutes of Health grants (DK118024 and DK125445 awarded to ECM, U01AI095473 awarded to GCH), the European Research Council ERC (694181), The Knut and Alice Wallenberg Foundation (2017.0028), Swedish Research Council (2017-00958) and the academy of medical sciences/Wellcome Trust through the springboard grant SBF005\1065 163470 awarded to AC. The authors acknowledge access to the SOLEIL and Diamond Light sources via both the University of Liverpool and Newcastle university BAGs (proposals mx21970 and mx18598, respectively). We thank the staff of DIAMOND, SOLEIL, and members of the Liverpool’s Molecular biophysics group for assistance with data collection. Mass spectrometry of glycans was performed in the Swedish infrastructure for biologic mass spectrometry (BioMS) supported by the Swedish Research Council. We are also grateful for Dr. Erwan Corre’s help regarding bioinformatics analyses (ABIMS platform, Station Biologique de Roscoff, France).

## Author contributions

ASL, AC and ECM designed experiments and wrote the manuscript. ASL and AC cloned, generated proteins mutants, expressed and purified sulfatases. ASL and AC performed enzyme assay, binding and kinetic analyses with assistance of DPB and PAE. MR and SO synthesized the O3-sulfated N-acetylgalactosamine. EAY labelled glycan substrates. AC grew, solved, collected and analysed crystallographic data with assistance from AB. AC and JAL carried out NMR kinetic analyses. CJ, ASL, GCH and NGK performed and interpreted analytical glycobiology experiments. ASL, GP, RWPG, SG, SS and NAP performed bacterial growth experiments and analysed *in vivo* competition data. MC, GM and TB performed sulfatase phylogenetic analyses. All authors read and approved the manuscript.

---

**Extended Data 1.**
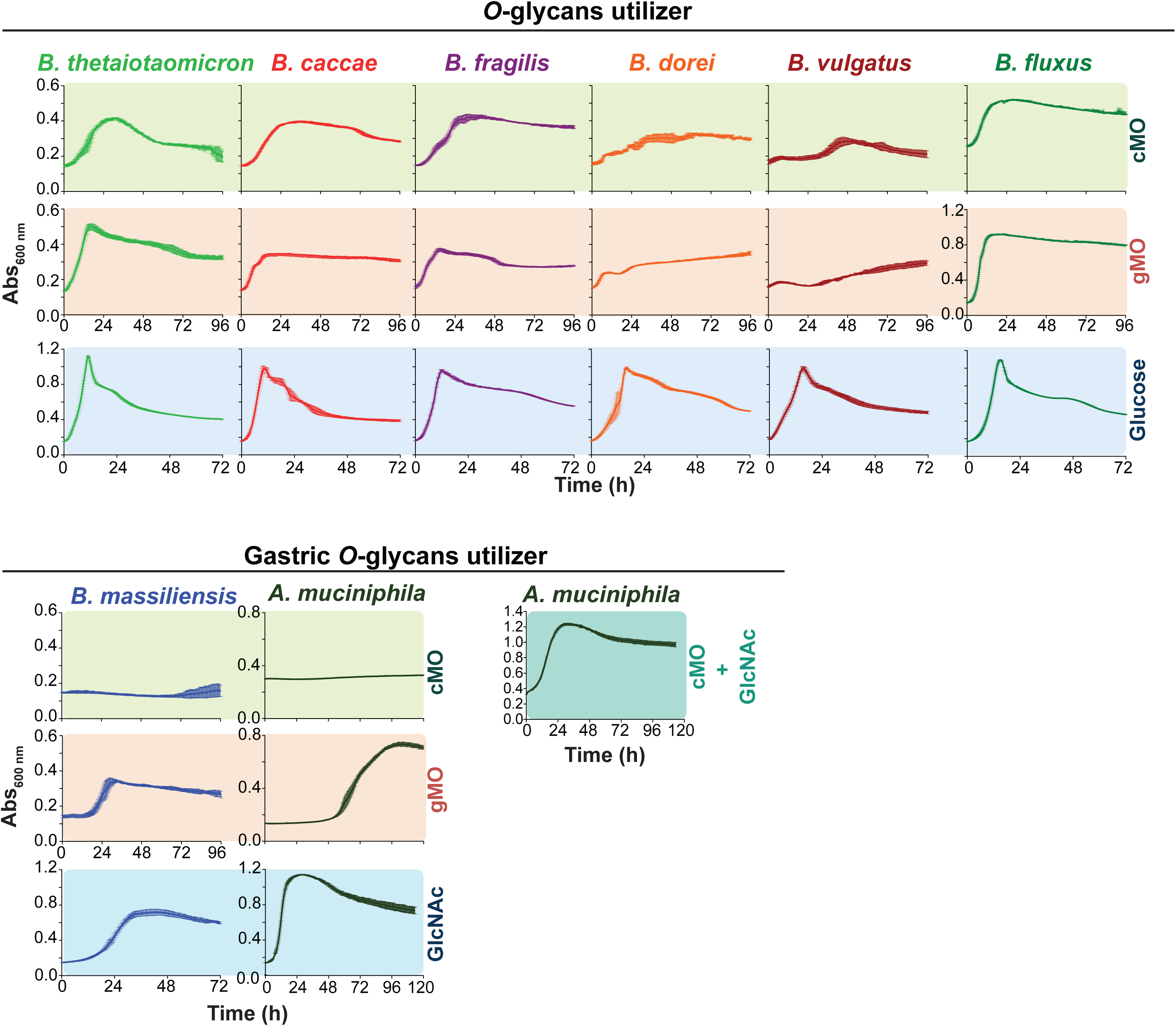
Growth of *Bacteroides* type strains and A*kkermansia muciniphila* in different mucin *O*-glycans. The graphics show the growth of strains able to utilize *O*-glycans in minimal media containing the indicated carbon source (biological replicates n =3, error bars denote the s.e.m.). cMO, colonic mucin *O*-glycans; gMO, gastric mucin *O*-glycans.

**Extended Data 2.**
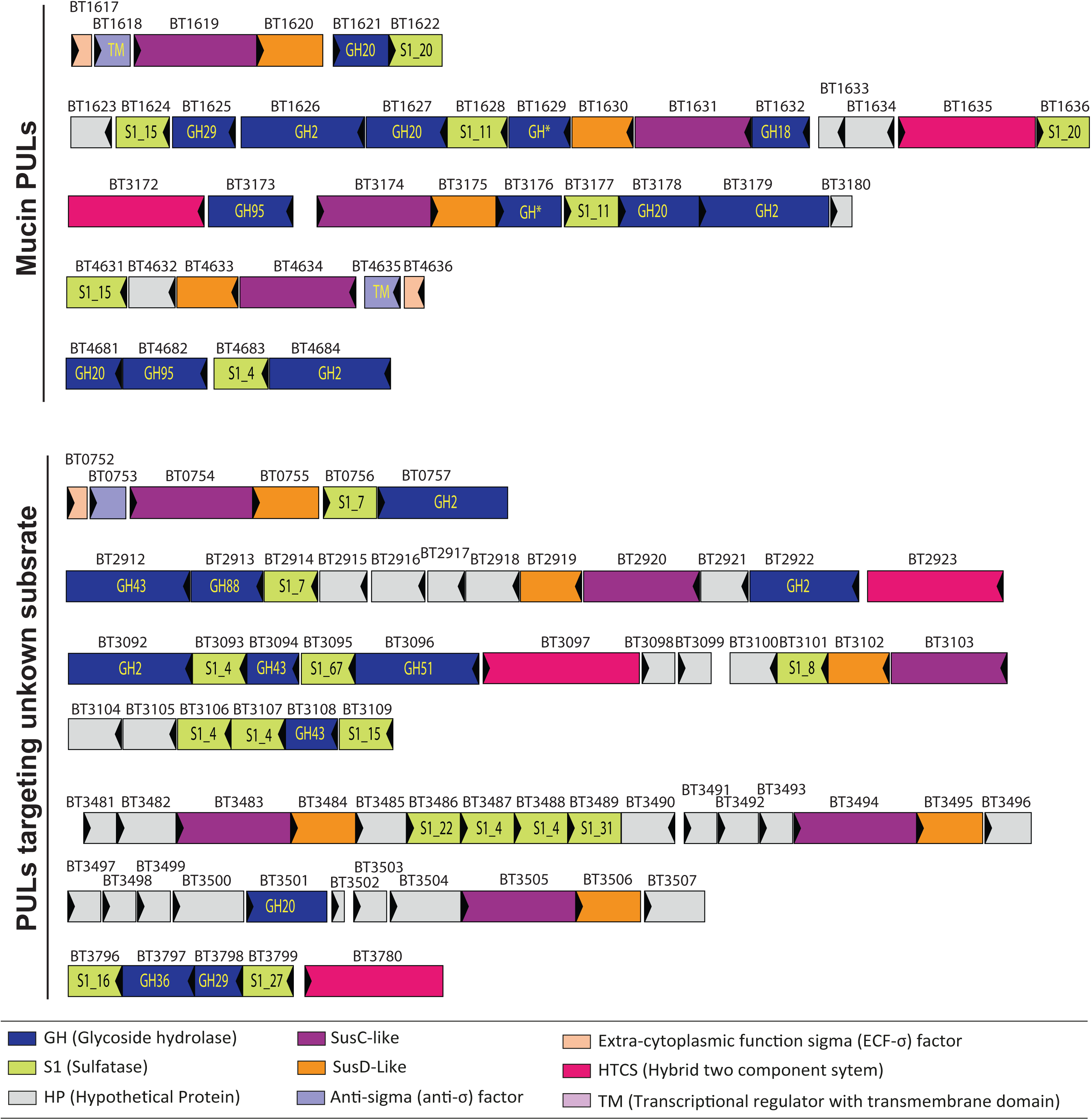
Schematic representation of polysaccharide utilization loci (PULs) encoding sulfatases (sulf). Genes are colour coded according to the predicted function of the respective proteins. Glycoside hydrolases (GH) in known families are indicated by GHXX or GH*, where XX and * indicates the respective family number or non-classified, respectively.

**Extended Data 3.**
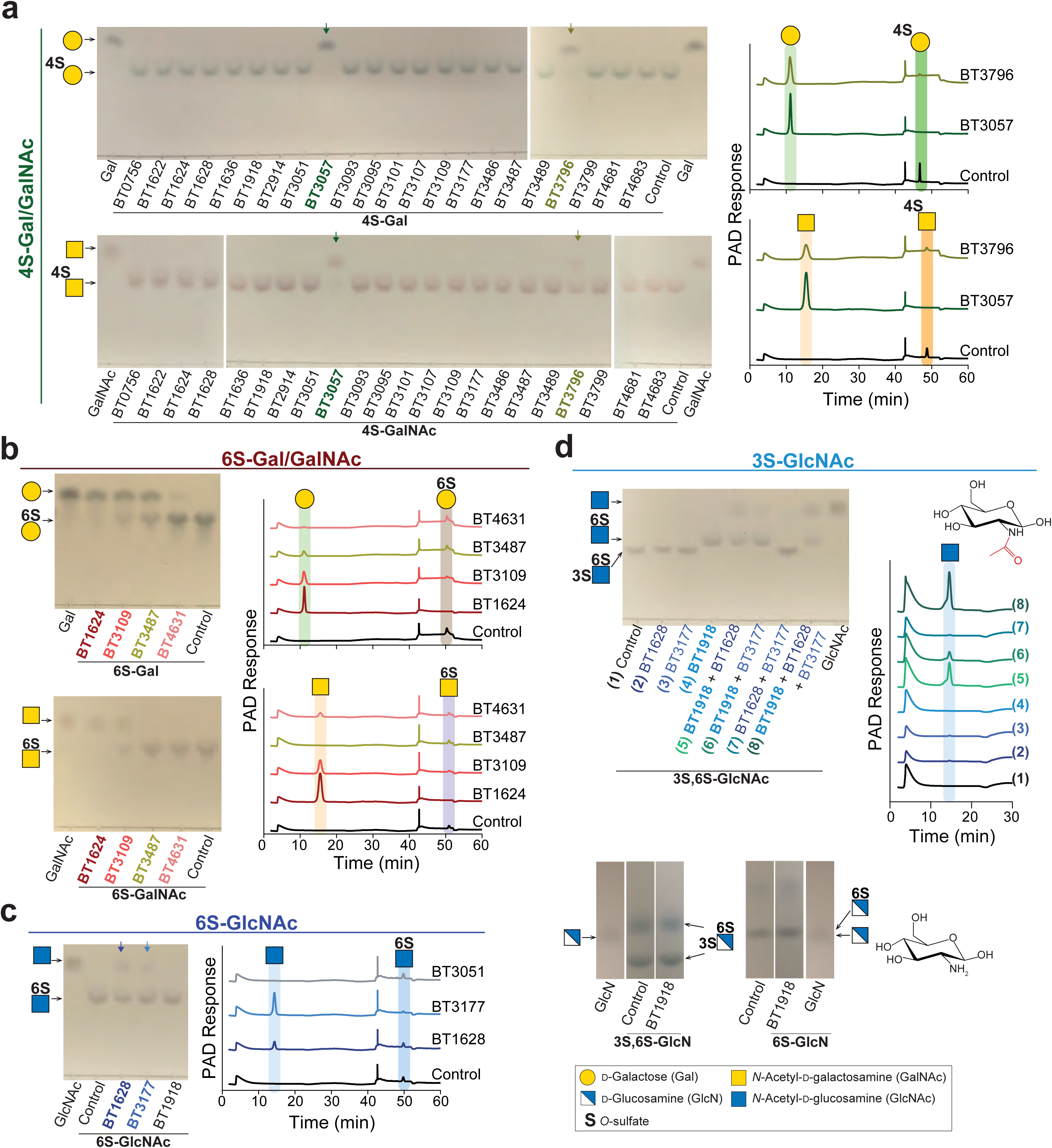
Enzymatic screen of *Bt* sulfatases using sulfated monosaccharides. Recombinant enzymes (1 *μ*M) were incubated with 1 mM of substrate in 10 mM MES pH6.5 with 5 mM CaCl_2_ for 16 h at 37°C. Reactions were analyzed by thin layer chromatography (left side) or HPAEC with pulsed amperometric detection (right side). Control reactions without sulfatases were carried out in the same conditions. The standards in TLC and HPAEC-PAD are labelled on the left side and top, respectively. The different panel represent activities found for sulfatases targeting: **(a)** 4S-Gal/GalNAc; **(b)** 6S-Gal/GalNAc; **(c)** 6S-GlcNAc and; **(d)** 3S-GlcNAc. The data shown are a representative from biological replicates (n = 3).

**Extended Data 4.**
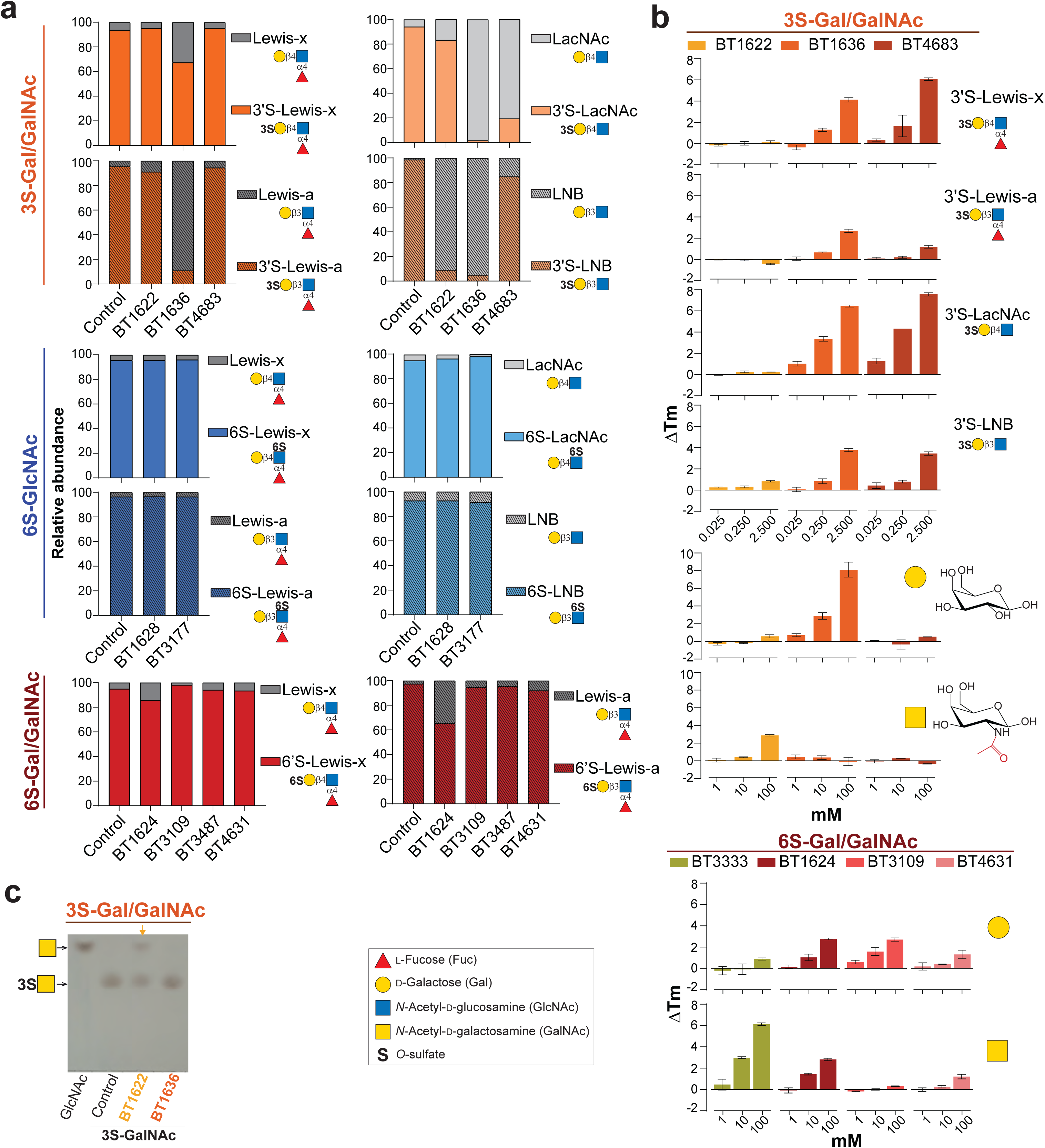
Activity and affinity of sulfatases to targeted substrates. **a,** Recombinant enzymes (1 *μ*M) were incubated with 1 mM of substrate in 10 mM MES pH6.5 with 5 mM CaCl_2_ for 16h at 37 °C. Sulfated disaccharides were generated by adding 1 *μ*M of a characterized *α*1,3/1,4-fucosidase (BT1625) in the enzymatic reaction. Control reactions without sulfatases were carried in the same conditions. Samples were analysed by mass spectrometry and the intensity of the substrate and reaction products was used for comparison of the relative abundance of these sugars after incubation with the respective enzymes. **b,** Affinity studies looking at the effect of ligand binding on the melting temperature of 3S and 6S-Gal sulfatases. All reactions were performed in 100 mM BTP, pH 7.0 with 150 mM NaCl. For sample melting temperatures see **Table S11**. **c,** Activity of 3S-Gal/GalNAc sulfatases (10 *μ*M) against 3S-GalNAc (10 mM). Reactions were performed in 10 mM Hepes, pH 7.0, with 150 mM NaCl and 5 mM CaCl_2_. The data shown are one representative from the biological replicates conducted (n = 3).

**Extended Data 5.**
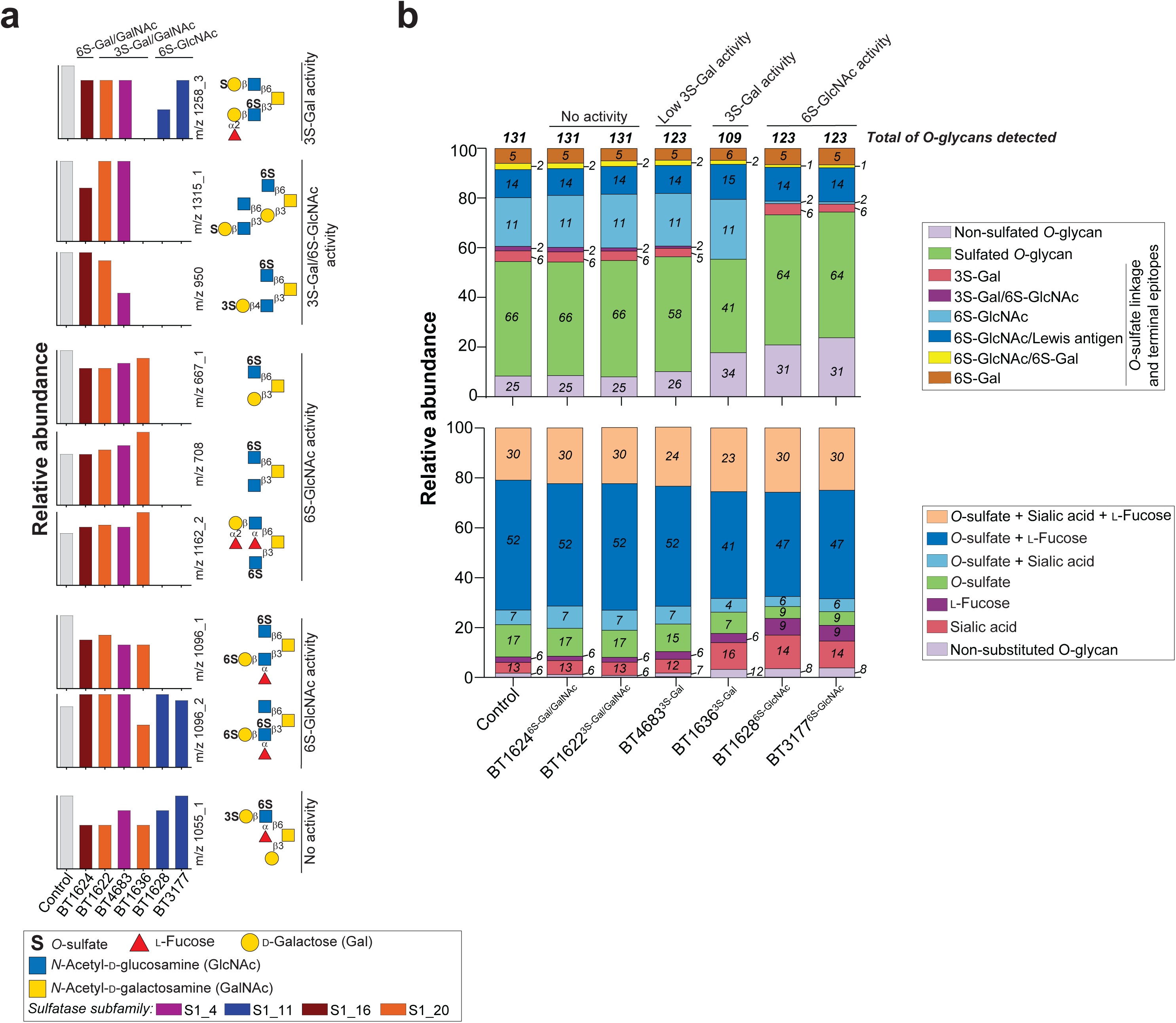
Activity of *Bt* sulfatases against colonic mucin *O*-glycans (cMO) analysed by mass spectrometry. **a,** Relative abundance of defined oligosaccharides after incubation of cMO with different sulfatases or no enzyme (control). The putative structure for the different mass is shown on the right side of the graphic. The reactions were performed with 1 *μ*M of enzyme and 0.5% cMO in 10 mM MES pH 6.5 with 5 mM CaCl_2_ for 16 h at 37°C. **b,** Relative abundance of structures detected in different samples organized by sulfate-linkage (top panel) or presence of one or several sugar substitutions such as sulfate, sialic acid and fucose (bottom panel). The colour-coded bars represent the relative abundance and the total number of the structures containing the specific linkage/substitution. The complete dataset is provided in **Supplementary Table 4**.

**Extended Data 6.**
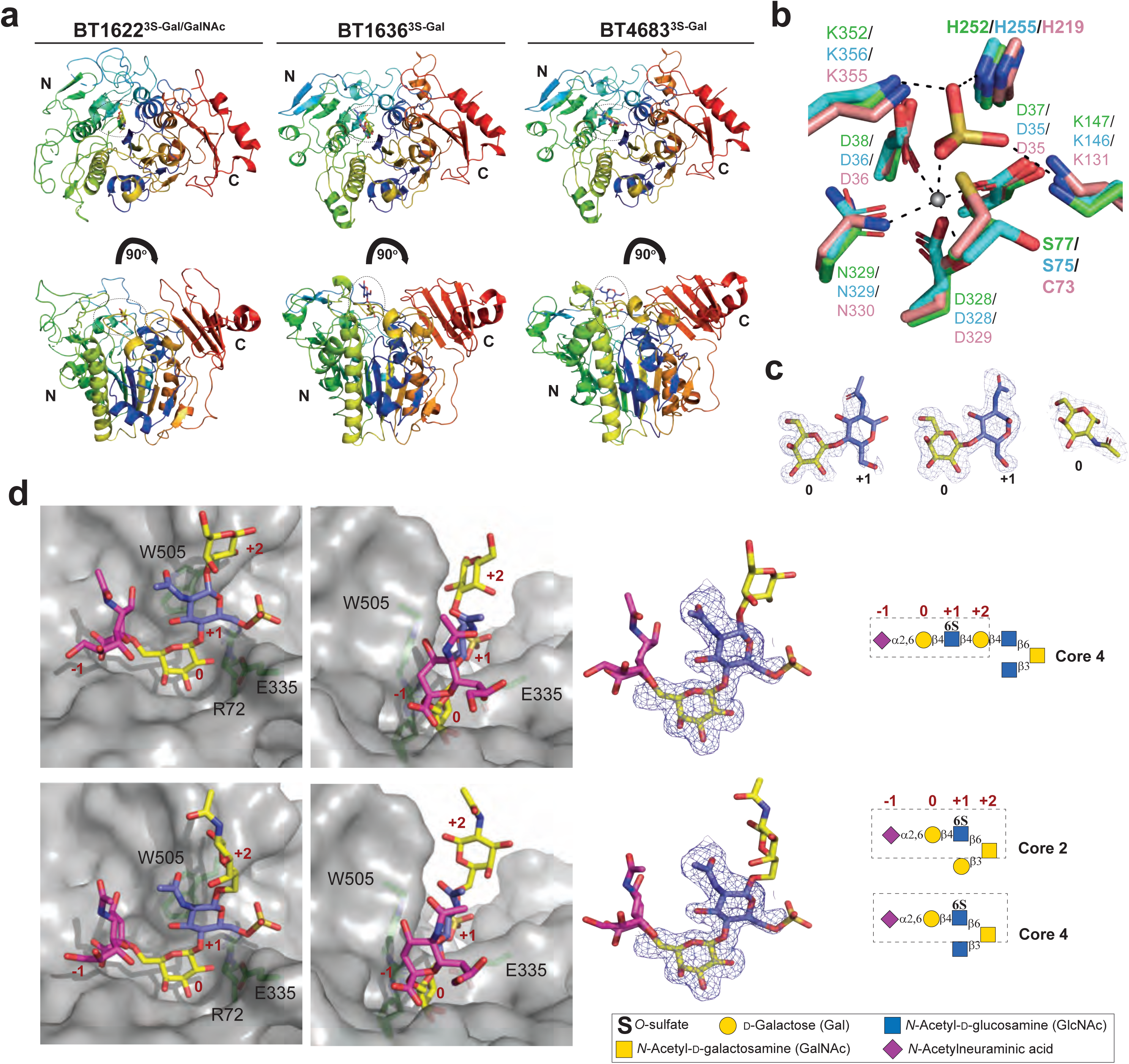
Schematic representation of 3S-Gal/GalNAc sulfatases. **a,** Cartoon representation colour ramped from blue (*α*/*β*/*α* N-terminal domain) to red (*β*-sheet C-terminal domain). **b**, Overlay of the active site S residues of BT1636^3S-Gal^ (green) BT1622^3S-Gal/GalNAc^ (blue) and BT4683^3S-Gal^ (pink). The putative catalytic residues are shown in bold. The calcium ion is represented as a grey sphere and its polar interactions indicated as dashed lines. **c,** Ligand density of maps for LacNAc in BT1636^3S-Gal^ and BT4683^3S-Gal^, and GalNAc in BT1622^3S-Gal/GalNAc^, contoured at 1*σ* (0.33 e/A^3^, 0.37 e/A^3^ and 0.18e/A^3^, respectively); **d,** Docking of putative structures of *O*-glycans targeted by BT4683^3S-Gal^ using the LacNAc as reference point showing that this structure can accommodate a sialic acid in -1 subsite and additional sugars in positive subsites (left hand side). The docking sugars are schematic shown as sticks (middle panel) and schematic represented inside the dashed box (right hand side). Using the LacNAc product as an ‘anchor’ additional sugars were built in manually with Coot 0.9 and regularized to low energy conformations.

**Extended Data 7.**
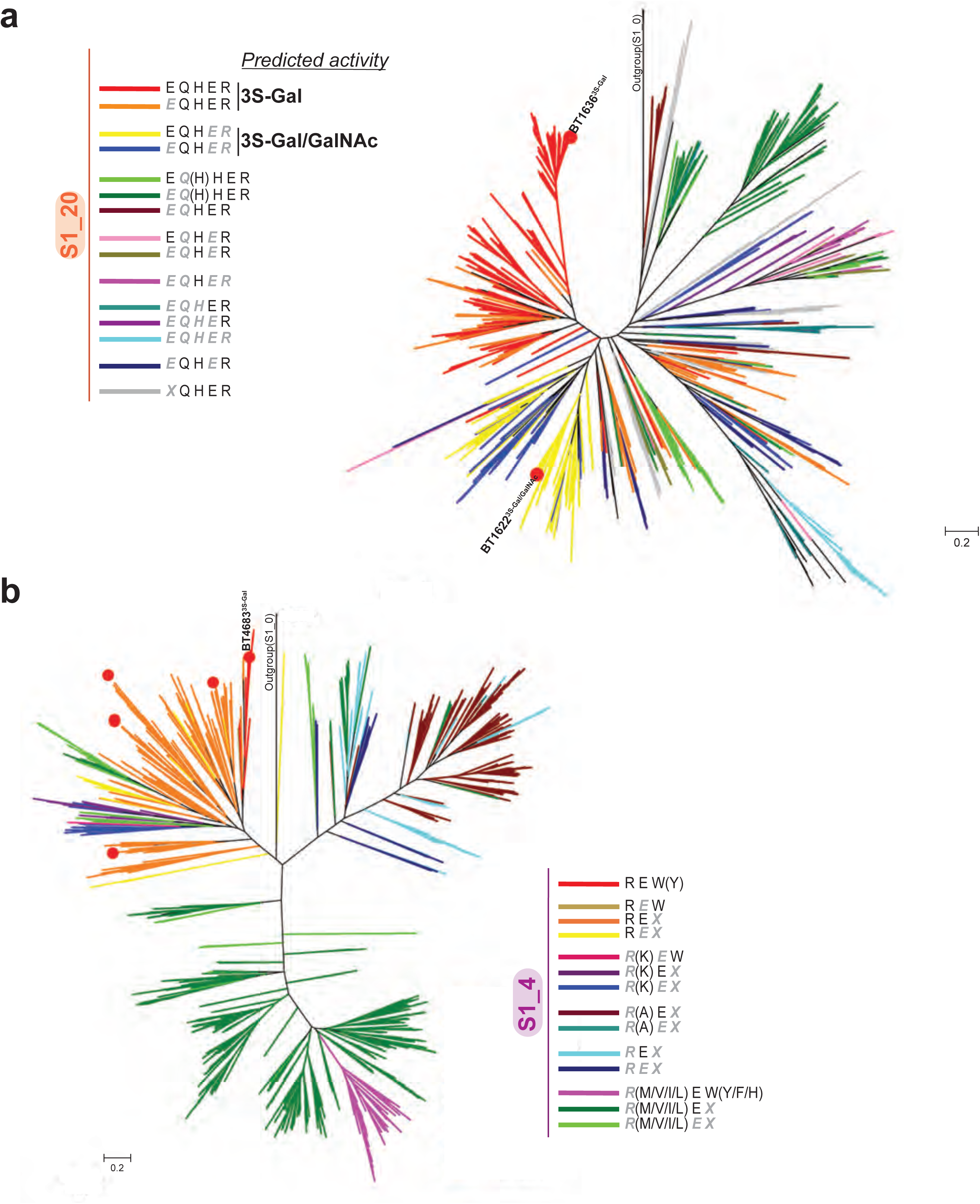
Phylogenetic tree of S1_20 and S1_4 sulfatases. The radial trees were constructed using the branched trees shown in **Supplementary Figs. 2 and 3**. For clarity, all labels and sequence accession codes have been omitted. Red filled circles designate sequences from *B. thetaiotaomicron* sulfatases. The residue is written in black without any attributes if present in the sequence, in grey and italics if the residue is mutated to any type in that sequence, or to a specific residue type if given in brackets. **a,** Radial representation of the phylogenetic tree constructed with representative sequences of the sulfatase S1_4 subfamily. The colour code is given as a pattern of presence or absence of the residues R72, E335 and W505, which are crucial in substrate recognition by BT4683 (acc-code Q89YP8, coloured red). A grey X in italics specifically designates that the residue W505 is absent in that sequence, and no obvious orthologous residue can be found from the alignment. **b,** Radial representation of the phylogenetic tree constructed with representative sequences of the sulfatase S1_20 subfamily. The colour code is given as a pattern of presence or absence of the residues E100, Q173 H177, E334, R353, which are crucial in substrate recognition by BT1636 (acc-code Q8A789, coloured red). A grey X in italics specifically designates that the residue E100 is absent in that sequence, and no obvious ortologous residue can be found from the alignment.

**Extended Data 8.**
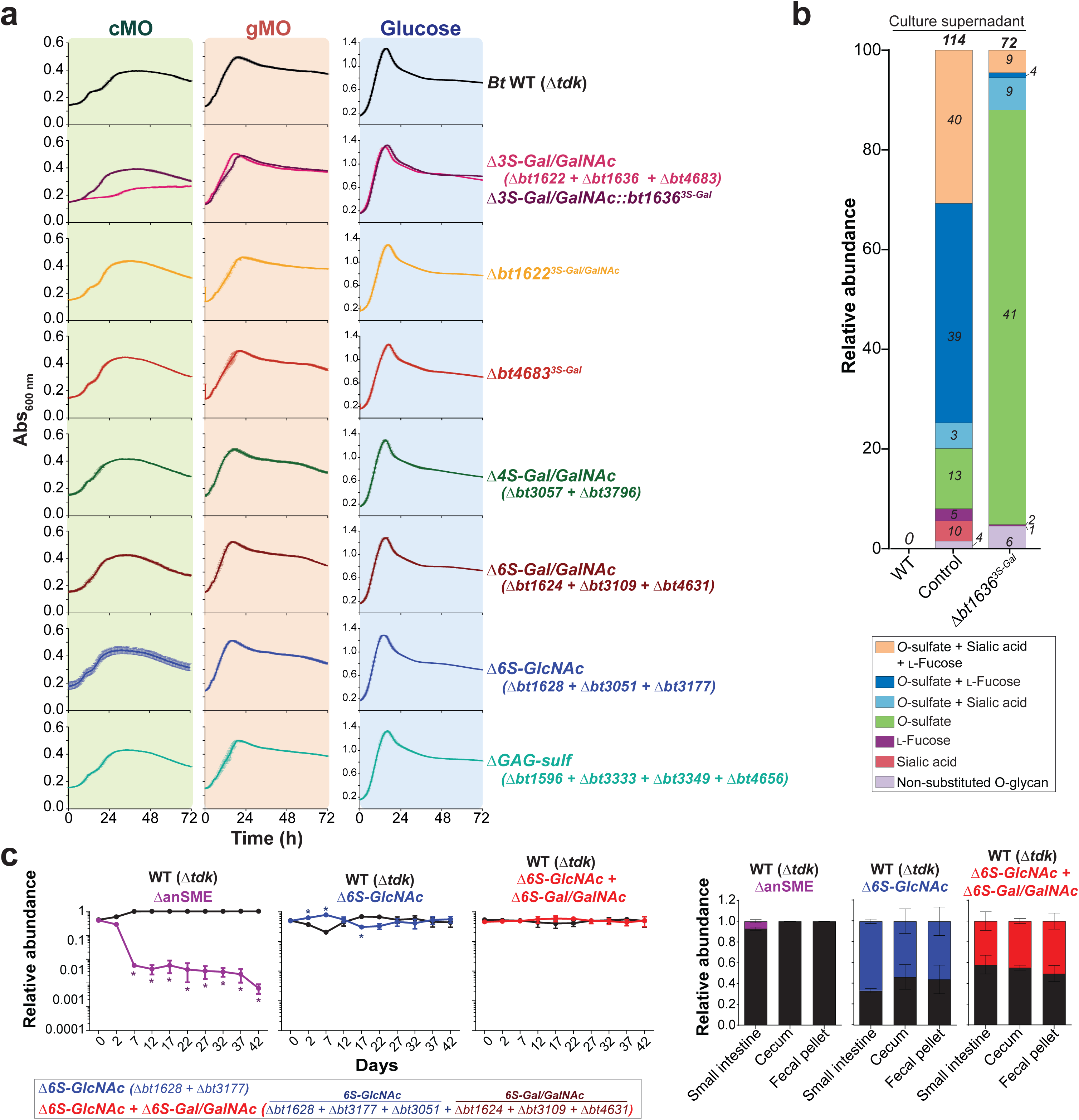
Sulfatase activity is required for growth in cMO and *in vivo* fitness. **a,** Growth curves of *Bt* wild-type *Δtdk* (WT), different sulfatase mutants (*ΔbtXXX*) and complemented strains on glucose, colonic or gastric mucin *O*-glycans (cMO and gMO, respectively). The curves represent the average of biological replicates (n = 3) and the error bars denote s.e.m. **b,** Relative abundance of oligosaccharides detected by mass spectrometry in culture supernatant of WT and *Δbt1636^3S-Gal^* after growth in cMO for 96h at anaerobic conditions. The control corresponds to cMO incubated in the same conditions without bacterium. The colours represent the relative abundance of structures grouped according to the presence of epitopes (sulfate, fucose and sialic acid) and the numbers represent the total number of structures that contain the respective substitution. **c**, Colonization of gnotobiotic mice fed a fiber-free diet by *Bt* WT and mutants lacking the full (*ΔanSME*, no S1 sulfatases active) or specific sulfatase activity (*Δ6S-GlcNAc* and *Δ6S-GlcNAc*+*Δ6S-Gal/GalNAc*). The fecal relative abundance of each strain was determined in regular intervals until day 42. The relative abundance of time 0 represents the abundance in gavaged inoculum. At the experimental endpoint the relative abundance was also determined in small intestine and cecum. The graphics represent the average of n =3 and the error bars denote the s.e.m. The relative abundance in each individual animal is represented in a lighter colour in each of the respective graphics.

## Supplemental Discussion

### 1. Utilization of different mucin O-glycans sources by HGM

Mucin composition varies throughout the gastrointestinal (GI) tract, with the stomach having mainly MUC5AC and the colon mainly MUC2^1^. The glycosylation of these respective mucins also varies along the GI tract with higher levels of sulfated and sialylated structures observed in the distal colon compared to the upper GI tract^2^. Among the 20 bacterial strains tested for growth, 12 failed to grow on gastric mucin *O*-glycans (gMO) or colonic mucin *O*-glycans (cMOs) (**Fig. 1b**). Only 6 bacteria were able to utilize both *O*-glycans substrates but growth was variable. In both *O*-glycan substrates, *Bacteroides thetaiotaomicron* (*Bt*), *B. caccae*, *B. fragilis* and *B. fluxus* grew better than *B. dorei* and B*. vulgatus* (**Fig. 1b** and **Extended Data Fig. 1**). The differences observed in the growth profiles were reproducible in two different batches of purified cMOs (**Fig. 1b**) Indeed, it is likely that different HGM members have evolved to target different (or only a subset) of the available *O*-glycans and this fine-tuning of host glycan utilization may have important implications in gut colonization and symbiosis. Additionally, *B. massiliensis* and *Akkermansia mucinipila* grew on gMO but failed to utilize cMO (**Fig. 1b** and **Extended Data Fig. 1**). Both strains were able to grow on *N*-acetylglucosamine (GlcNAc) and *Akkermansia mucinipila* grew on GlcNAc in the presence of cMO suggesting that these *O*-glycans do not inhibit the growth of this bacterium. Previous studies have determined that *B. massiliensis* and *Akkermansia mucinipila* are mucin-degraders by demonstrating growth on gastric mucins^3, 4^. However, the lack of growth in colonic *O*-glycans suggests that these bacteria are not able to initiate the degradation of more complex, sulfated colonic glycans. This finding highlights the importance of taking into account *O*-glycosylation differences along the GI tract and the need to utilize colonic mucins to draw conclusions regarding the full mucin-degrading potential of the colonic HGM.

### 2. Sulfatase activity in cMO

Despite all 12 sulfatases being active on defined oligosaccharides, of those tested on cMO, BT1622^3S-Gal/GalNAc^ (S1_20 subfamily) and BT1624^6S-Gal/GalNAc^ (S1_15 subfamily) did not show any activity on this complex substrate (**Fig. 2, Extended Data Fig. 5** and **Supplementary Table 4)**. These findings are consistent with the results observed in defined commercial oligosaccharides where BT1622^3S-Gal/GalNAc^ showed a preference for sulfated GalNAc over Gal glycans (**Extended Data Fig. 4**) and BT1624^6S-Gal/GalNAc^ activity is blocked by the presence of additional substitutions (such as Lewis antigens) (**Extended Data Fig. 4a**). Additionally, 2 of 6 detected 6S-Gal structures contained a capping sialic acid and a terminal blood group H type 2 [Fuc-*α*1,2-(6S)Gal-*β*1,4-GlcNAc-] (**Fig. 2**). The lack of activity of BT1624^6S-Gal/GalNAc^ towards such structures confirms an exo-mode of action that we describe for this sulfatase using commercial substrates.

Overall, when compared to the non-enzyme treated control, we detected an increase of non-sulfated structures and decrease of sulfated oligosaccharides in all samples with the active enzymes (BT4683^3S-Gal^, BT1636^3S-Gal^, BT1628^6S-GlcNAc^ and BT3177^6S-GlcNAc^) (**Extended Data Fig. 5b**). BT1636^3S-Gal^ (S1_20) was active towards all detected 3S-Gal structures with the exception of glycan 1055_1 that is a doubly sulfated 3S-Gal/6S-GlcNAc fucosylated structure (**Extended Data Fig. 5a**). As we observed using commercial substrates, the presence of Lewis-a/x epitopes leads to a decrease in the activity of this sulfatase (**Extended Data Fig. 4a** and **Supplementary Table 4**) and the presence of a second sulfate group might exacerbate this negative effect leading to the lack of activity towards this complex sulfated *O*-glycan. The incubation of the 6S-GlcNAc sulfatases BT1628^6S-GlcNAc^ and BT3177^6S-GlcNAc^ with cMO suggests that these enzymes are redundant, but because they are encoded in different PULs they could be expressed in response to different activating cues (**Fig. 2** and **Supplementary Table 4)**. Compared to the non-enzyme treated control, 16 glycans were not detected after incubation with these sulfatases, 14 of these structures have a terminal 6S-GlcNAc (**Fig. 2** and **Supplementary Table 4)**. BT1628^6S-GlcNAc^ and BT3177^6S-GlcNAc^ were active in 6S-GlcNAc core 3 (GlcNAc-*β*1,3-GalNAc) and core 4 (GlcNAc-*β*1,6-GalNAc) structures (**Extended Data Fig. 5a**), suggesting that these sulfatases are well suited to accommodate the variations in linkages/sugars found in mucin *O*-glycans. Additionally, we also detect 7 new glycans that are likely to be reaction products of BT1628^6S-GlcNAc^ and BT3177^6S-GlcNAc^ (**Fig. 2** and **Supplementary Table 4)**.

The identification and characterization of the first sulfatases active on mucin *O*-glycans creates the opportunity to improve our understanding of *O*-glycan structures by using these enzymes as analytical tools. After the treatment with BT1636^3S-Gal^ several oligosaccharides predicted to contain a terminal sulfate linked to Gal were not detected. Although we could not determine the specific sulfate linkage by mass spectrometry, the activity of the 3S-sulfatase suggests that these oligos contain a terminal 3S-Gal (**Extended Data Fig. 5a** and **Supplementary Table 4**). The specificity of the 6S-GlcNAc sulfatases for non-fucosylated *O*-glycans also illuminates their potential use as tools to characterize the structure of these complex structures since it allows the differentiation of different isomers. For example, we detect two oligosaccharides with mass 1096, however, after incubation with BT1628^6S-GlcNAc^ or BT3177^6S-GlcNAc^, only the isomer 1096_2 was detected, indicating that the isomer 1096_1 contain a terminal 6S-GlcNAc (**Extended Data Fig. 5a** and **Supplementary Table 4**).

### 3. Conserved structural features of the S1 formylglycine family

#### Protein fold and subsites nomenclature

S1 sulfatases comprise the most common and largest family of sulfatases, currently encompass 36,816 members in sulfAtlas and are found in all domains of life^5^. S1 sulfatases are part of the alkaline phosphatase superfamily and adopt an alkaline phosphatase-like fold. This is an N-terminal *α*/*β/a* domain with S1 sulfatases also possessing a smaller C-terminal ‘sub domain’. The active site is located in the N-terminal domain that has a large mixed *β*-sheet composed of ∼10 *β* strands, sandwiched between *α* helices above and below. The C-terminal ‘sub-domain’ is composed of a 4 stranded antiparallel *β*-sheet and a single amphipathic terminal helix. This C-terminal domain abuts the N-terminal domain through the antiparallel *β*-sheet with loops from the *β* strands sometimes contributing to the active site architecture (**Extended Data Fig. 6a**). The subsite nomenclature for carbohydrate sulfatases is such that the invariant sulfate binding site is denoted as the S site. The S site sulfate is appended to the 0 subsite sugar. Subsites then increase in number (i.e. +1, +2, +3) as the sugar moves toward the reducing end (free O1) and decreases in number as the sugar chain moves towards the non-reducing end (i.e. -1, -2, -3)^6^.

#### S1 formylglycine active site conservation

The sulfate binding site (S site) is invariant across the S1 family and comprises the catalytic residues (nucleophile and catalytic acid) and a calcium binding site (**Extended Data Fig. 6b**). An invariant histidine is likely the potential catalytic acid but a lysine has also been suggested to possibly fulfil this role^7^. The pKa of His is ∼6.0, whilst Lys has a pKa of >10, making it more chemically feasible that His performs the role of the catalytic acid. Homologues of these residues (H252 and K352 in BT1636^3S-Gal^) make hydrogen bonds to the scissile sulfoester linkage (**Extended Data Fig. 6b**). Previously published work with BT1596 and BT4656, which are 2S-Uronic acid and 6S-GlcNAc sulfatases, respectively, showed that the mutation of either residue to alanine causes inactivation^7^. Consistent with this work, a BT4683^3S-Gal^ H219A mutant was inactive. However, in BT1622^3S-Gal/GalNAc^, the mutation of H255 to Ala caused only a ∼30-fold decrease in activity (**Supplementary Table 3**). Thus, it is possible that in BT1622^3S-Gal/GalNAc^ the loss of H255 is compensated by the invariant residue K356 and interestingly BT1622^3S-Gal/GalNAc^ has a pH optimum ∼2 units higher than most sulfatases assayed (**Supplementary Fig. 4**).

The calcium binding site is located at the base of the S site interacting with the sulfate group. This calcium ion is an essential component of the catalytic mechanism helping to stabilise negative charges that occur during the catalysis. All three of the solved structures had occupation for calcium. In BT1636^3S-Gal^ D328 and the sulfate group of the substrate coordinate above and below the calcium with D37, D38, N329 and the formylglycine binding in a plane completing an octahedral coordination (**Extended Data Fig. 6b**). These three Asp and the Asn coordinated with calcium are structurally conserved in all 3S-Gal/GalNAc sulfatases structures (**Extended Data Fig. 6b**).

The solved structures of BT1636^3S-Gal^ and BT1622^3S-Gal/GalNAc^ were native *Bt* proteins having a Ser at the formylglycine position. However, the structure of BT4683^3S-Gal^ was obtained with the active protein where S73 was mutated to Cys (as *E. coli* can only convert Cys, not Ser, to formylglycine). The analysis of BT4683^3S-Gal^ reveals that the crystallized protein still has the Cys and not formyglycine indicating poor installation of the formylglycine. This observation means the kinetic data, (although the rates are significant and readily measurable) may be an underestimation of true catalytic performance. This will affect the *k*_cat_ component of the *k*_cat_/*K*_M_ measurement and thus the *k*_cat_/*K*_M_ reported in **Supplementary Table 3** is an underestimate of the true activity.

#### Additional 3S-Gal/GalNAc specificity determinants based on structures

BT1636^3S-Gal^ was solved in complex with the product LacNAc; the Gal at 0 subsite is well ordered and makes extensive interactions, whilst the +1 GlcNAc is highly disordered and appears to make no interactions with the protein (**Fig. 3** and **Extended Data Fig. 6c**). O2 of the Gal hydrogen bonds with O*ε*1 of E334 and NH2 of R353. Mutation of these residues to Ala causes ∼300 and ∼60-fold reductions in *k*_cat_/*K*_M,_ respectively. The O6 group of Gal potentially coordinates with O*ε*2 of E100 and N*ε*2 of Q173 and mutations of these residues to Ala cause ∼80 and ∼50-fold decreases in *k*_cat_/*K*_M,_ respectively (**Fig. 3** and **Supplementary Table 3**). Comparison of the BT1622^3S-Gal/GaNAc^ structure with BT1636^3S-Gal^ shows that E98 and Q172 (which correspond to E100 and Q173 in BT1636^3S-Gal^) are conserved **(Fig. 3)** and mutating E98 to Ala caused only a 15-fold decrease in *k*_cat_/*K*_M_ (**Supplementary Table 3**). Additionally, in BT1622^3S-Gal/GaNAc^ the hydrophobic interactions with the *N*-acetyl group, and the more open pocket, may offset the effects H176A (300-fold loss in activity) when compared to H177A (complete loss in activity) in BT1636^3S-Gal^ (**Fig. 3** and **Supplementary Table 3**).

BT4683^Gal-3S^ also displayed the same 3S-Gal activity as the S1_20 enzymes but showed a preference for 3’S-LacNAc, reciprocal to BT1622^3S-Gal^. BT4683^3S-Gal^ bound the O2 Gal of LacNAc via O*ε*2 of E335 (equivalent to E334 in BT1636^3S-Gal^) and through either N*ε* or NH1 of R72 **(Fig. 3)**. Although R72 is sequentially distal to R353 in BT1636^3S-Gal^ it is spatially similar and likely contributes in a similar capacity **(Fig. 3)**. Despite the mutations R72A and E335A resulting in loss of activity, the Glu and Arg are only conserved in 62 % and 19 % of S1_4 sequences, respectively, suggesting there is a significant but not absolute selection for an equatorial O2 in this subfamily (**Extended Data Fig. 7** and **Supplementary Fig. 2**). Uniquely among the 3S-Gal sulfatases identified, BT4683^3S-Gal^ utilises a hydrophobic stacking interaction through W505 to provide a platform for the +1 GlcNAc and partially the 0 Gal. Mutation of W505 to Ala almost completely abolishes activity on 3’S-LacNAc (**Supplementary Table 3**) but surprisingly this residue is not conserved in our phylogenetic analyses of S1_4 being present in only 8 other sequences (**Extended Data 7** and **Supplementary Fig. 2**). It is important to note, that W505 is not well conserved; potential equivalent aromatic residues can be found in some additional clades, which are coloured light brown (or bronze), pink or dark red, but it is not evident from the alignment that these are functional equivalents. Future structural work is needed to confirm if other aromatic residues take equivalent positions in those sulfatases. Additionally, the BT4683^3S-Gal^ activity against defined sulfated saccharides was suggestive of an exo-acting enzyme that cleaves terminal 3S-Gal (**Extended Data Fig. 4**). However, a close analysis of this sulfatase structure shows that the active site is located in an open cleft characteristic of an endo-active enzyme^6^. This more open cleft of BT4683^3S-Gal^ may allow additional sugars/sulfates to be accommodated on the O6 of both the 0 Gal and +1 GlcNAc. Indeed, the activity determined in cMO shows that this sulfatase can act on sialylated *O*-glycan (**Fig. 2**). Further modelling of different *O*-glycan structures (using the crystallographically solved LacNAc as an ‘anchor’) indicate that this enzyme can accommodate complex *O*-glycans with internal sulfation **(Fig. 3 and Extended Data 6d)**. Together, these results suggest that BT4683^Gal-3S^, and its close homologues, could be endo 3S sulfatases where the 0 subsite specificity for Gal is driven by glycan context and/or distal subsites such as -1 and +2, rather than an axial O4 as in S1_20.

Additionally, it is unclear why BT1636^Gal-3S^ acts better on LacNAc substrates than BT1622^3S-Gal/GalNAc^. It is interesting to note, however, that both BT1636^3S-Gal^ and BT4683^3S-Gal^ perform well on LacNAc configured substrates and utilise an Arg and Glu to coordinate O2 whilst BT1622^3S-Gal^ lacks these residues **(Fig. 3)**. These residues may lead to the enhanced activity on LacNAc (*β*1,4 glycan) vs. LNB (*β*1,3 substrate). Another thing to note is that a *β*1,4 vs *β*1,3 linkage will rotate the GlcNAc ∼60° but switch the position of the Fuc residue from being on the ‘*N*-acetyl side’ of the glycosidic bond to the ‘O6 side’ of the glycosidic bond, and this may also be the cause of the differential activities on *β*1,4 vs *β*1,3 linked substrates.

#### Phylogenetic analyses of S1_20 specificity determinants

The essential His that acts as a key specificity determinant of galacto-over gluco-substrates (H177 and H176 in BT1636^3S-Gal^ and BT1622^3S-Gal/GalNAc^, respectively) is highly conserved (92% of S1_20 sequences) (**Extended Data Fig. 7** and **Supplementary Fig. 3**). The Gln (Q173 and Q172 in BT1636^3S-Gal^ and BT1622^3S-Gal/GalNAc^, respectively) is only conserved in 66% of sequences and in 25% of the cases is substituted with a histidine, a residue that can also fulfill the same role of Gln interacting with Gal O6. Indeed, these conserved residues are located in a highly conserved domain with the consensus sequence [CDNS]-[QH]-[RVF]-[QHLD]-[AG]-H-[NRST]-[YHF]-[YF]-P (Prosite syntax). With H177 targeting the axial O4 of Gal directly, a Q173 may function indirectly to select for an axial O4 and thus these residues may operate as a selectivity ‘dyad’ for Gal with S1_20. Additionally, the residues implicated in recognition of Gal over GalNAc, E335 and R353 in BT1636^3S-Gal^ are conserved in 64 and 74% of S1_20 sequences, whilst the residue that allows the accommodation of O2 N-acetyl and activity in GalNAc (N334 in BT1622^3S-Gal/GalNAc^) is only found in 8% of members of this family (**Extended Data Fig. 7** and **Supplementary Fig. 3**). This observation suggests that the majority of the S1_20 sulfatases evolved to target sulfated Gal and only a subset of this subfamily’s members can actually also be active on GalNAc. Interestingly, all of the close homologs of BT1636^3S-Gal^ and BT1622^3S-Gal/GalNAc^ that share the critical specificity determinants of these proteins (**Supplementary Tables 12 and 13**) were isolated from mammals at body regions rich in mucins, highlighting the role of these sulfatases in accessing sulfated host glycans.

### 4. Growth of sulfatase mutants on *O*-glycans

The deletion strain lacking 4S-Gal/GalNAc sulfatases (*Δbt3057 + Δbt3796*) did not show any phenotype in cMO (**Extended Data Fig. 8a**), a result that is consistent with the lack of these sulfated linkages in colonic mucins (**Supplementary Table 4**). Unexpectedly, the deletion strains lacking the identified 6S-Gal/GalNAc sulfatases (*Δbt1624 + Δbt3109 + Δbt4631*) and 6S-GlcNAc sulfatases (*Δbt1628 + Δbt3051 + Δbt3177*) also did not show any growth defect on cMOs (**Extended Data Fig. 8a**). Analysis of cMO by mass spectrometry showed that this substrate contains a low abundance of 6S-Gal but a relatively high abundance of 6S-GlcNAc, especially in shorter structures (**Supplementary Table 4**). Although the low abundance of O6- sulfated Gal could explain the lack of phenotype of the 6S-Gal/GalNAc sulfatase deficient strain, the lack of effect in the *Δ6S-GlcNAc* mutant in cMO was unexpected (**Extended Data Fig. 8a).** Due to the limitations of the mass spectrometry technique it is not possible to analyse sulfation in longer oligos, making the real complexity of glycans found in colonic mucins unclear. Indeed, the lack of phenotype of *Δ6S-GlcNAc* mutant in cMO suggests that 6S-GlcNAc might not be a major terminal epitope in colonic mucins. It is also important to note that the mutant *Δ6S-GlcNAc* is the deletion of two characterized 6S-GlcNAc sulfatases active on cMO (BT1628^6S-GlcNAc^ and BT3177^6S-GlcNAc^) and a third closely related S1_11 sulfatase (BT3051^putative_6S-GlcNAc^) for which no activity was found. This putative 6S-GlcNAc sulfatase was deleted to avoid possible compensation of function after loss of BT1628^6S-GlcNAc^ and BT3177^6S-GlcNAc^ activities.

The deletion of previously characterized GAG-specific sulfatases^8^ (*Δbt1596 + Δbt3333 + Δbt3349 + Δbt4656*) did not result in any observable phenotype in cMO (**Extended Data Fig. 8a**), indicating that this substrate was not contaminated with additional endogenous host glycans. Additionally, despite some mutants exhibiting growth defects on sulfated cMO, all of the mutants grew well on gMO and glucose (**Fig. 4a** and **Extended Data Fig. 8a**), suggesting that the phenotypes observed are dependent on the mucin source (colon) and cannot be observed utilizing mucins from other regions of the gastrointestinal tract. Together these results highlight the contribution of sulfatases in utilization of colonic mucins by the HGM.

### 5. Analysis of Δ*bt1636^3S-Gal^* culture supernatant by MS

The analysis of the oligosaccharides present in *Δbt1636^3S-Gal^* culture supernatant after 96h incubation revealed that the detected glycans are different from the cMO profile in the starting material (**Fig. 4c**, **Extended Data Fig. 8b** and **Supplementary Table 5**). We detected 114 glycans in the cMO sample, of which 39 were sulfated and fucosylated (44% total) (**Extended Data Fig. 8b**) and the three most common structures (12% total) were 6S-GlcNAc oligosaccharides (**Fig. 4c** and **Supplementary Table 5**). In the control sample, the levels of sulfation, sialylation and fucosylation were 92%, 40% and 77%, respectively **(Supplementary Table 5**). In the *Δbt1636^3S-Gal^* culture supernatant, we detected 72 glycans, of which 41 were substituted only with *O*-sulfate (84% total) (**Extended Data Fig. 8b**). In the mutant supernatant the levels of sulfation (95%) were similar to cMO, however the levels of sialylation (11%) and fucosylation (5%) decreased substantially (**Supplementary Table 5**), suggesting that this mutant is not able to utilize sulfated structures and these accumulate in culture media.

Additionally, a total of 98 of the 114 structures present in cMO were not detected in *Δbt1636^3S-Gal^* culture supernatant whereas in mutant supernatant, we detected 49 glycans that were not detected in the initial substrate (**Supplementary Table 5**). This suggests that some of the oligosaccharides present in cMO can support the limited growth of *Δbt1636^3S-Gal^* and, although this mutant is not able to utilize many sulfated cMO structures, it can still modify the glycans to create novel structures. It remains unclear which enzymes are encoded by the mutant to modify the *O*-glycans, but the presence of a cell surface sialidase^9^ can explain the decrease of sialylation levels in structures found in *Δbt1636^3S-Gal^* supernatant. Additionally, the presence of surface endo-acting glycoside hydrolases able to cleave *O*-glycans into shorter oligosaccharides^10^ can also contribute to new glycan structures in the mutant culture supernatant. Together these results show that *Δbt1636^3S-Gal^* is not able to utilize most sulfated *O*-glycans explaining the limited growth of this mutant in cMO.

---

**Supplementary Table 1.**
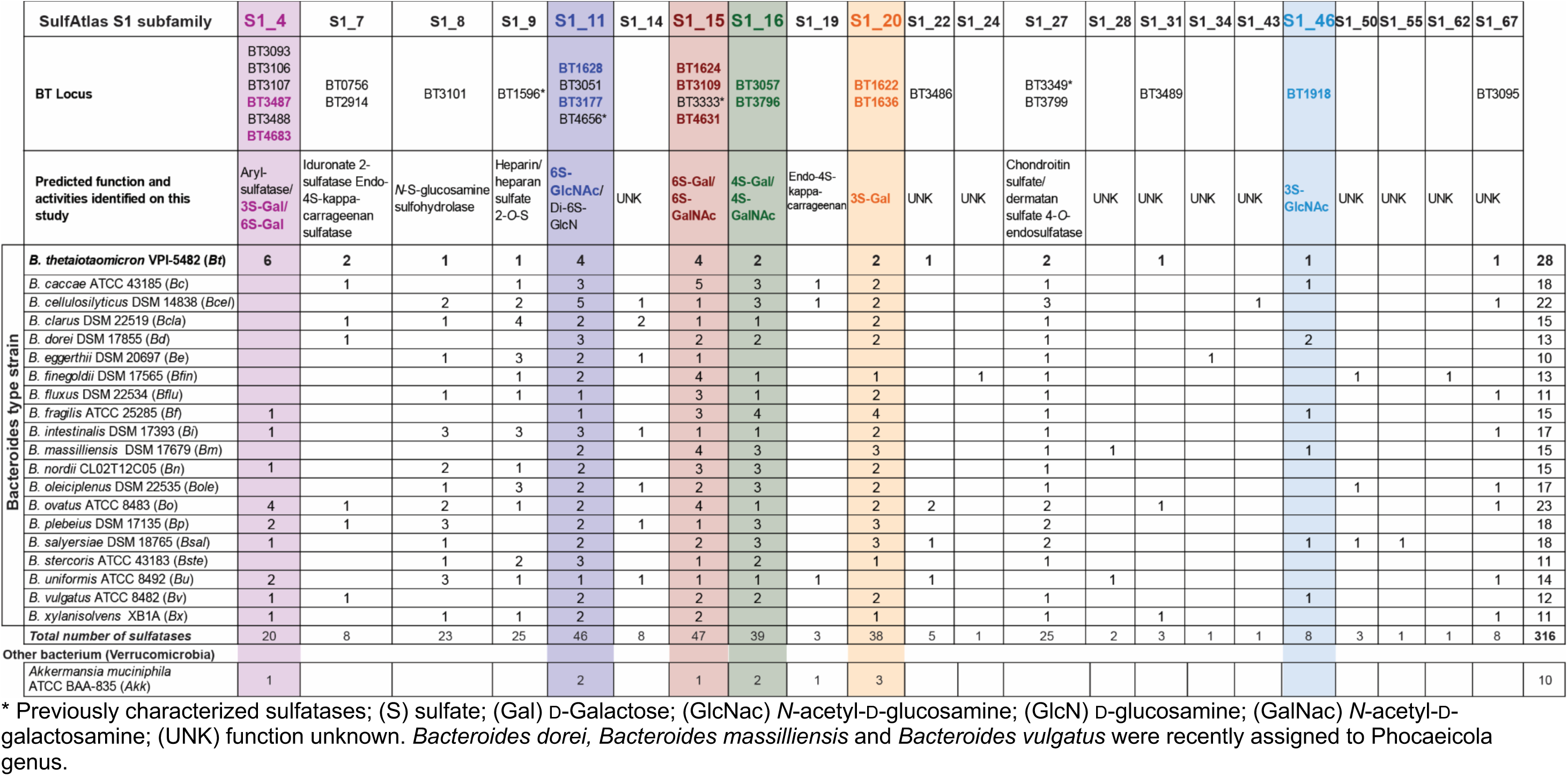
Family S1 sulfatase subfamiles encoded in the genomes of different *Bacteroides* type strains and *Akkermansia*

**Supplementary Table 2.**
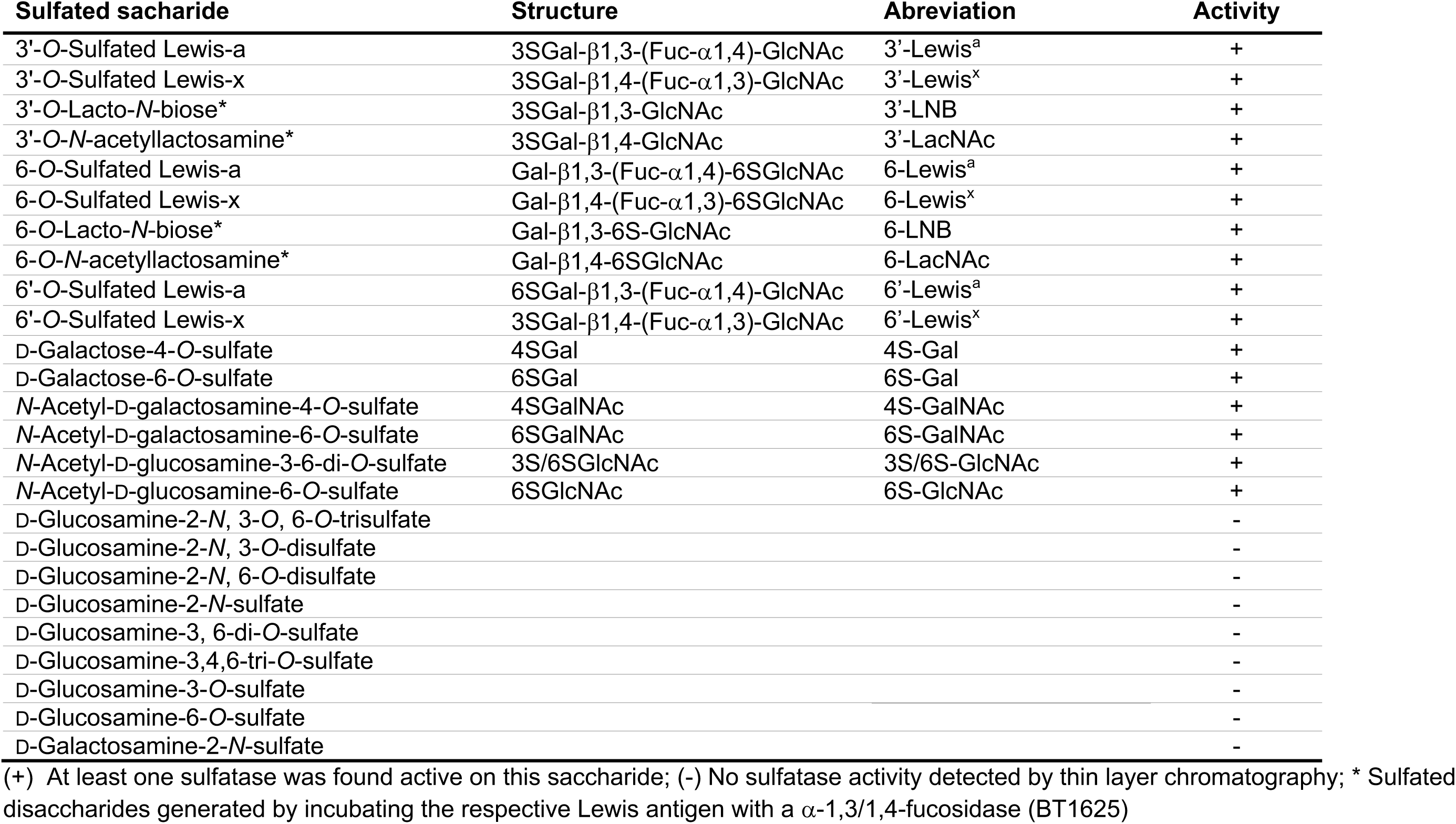
List of sulfated saccharides used in the initial sulfatase activity screen

**Supplementary Table 3.**
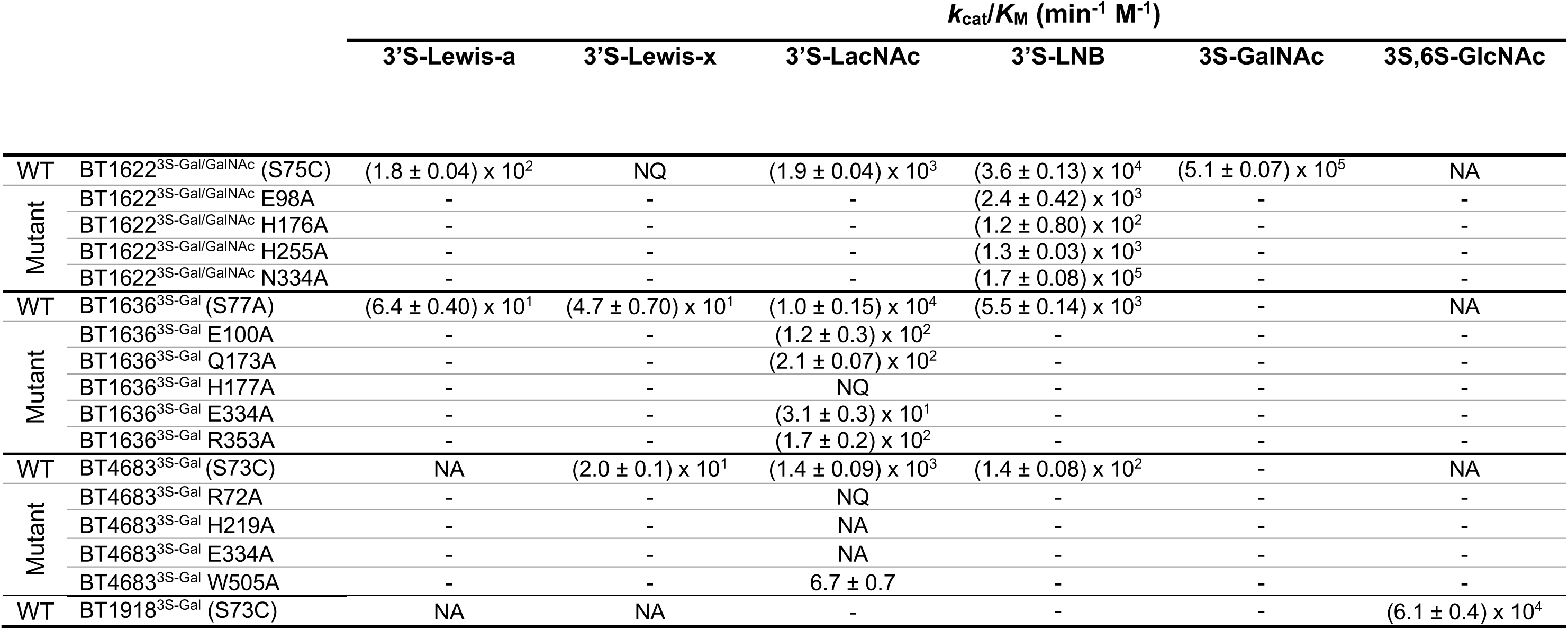
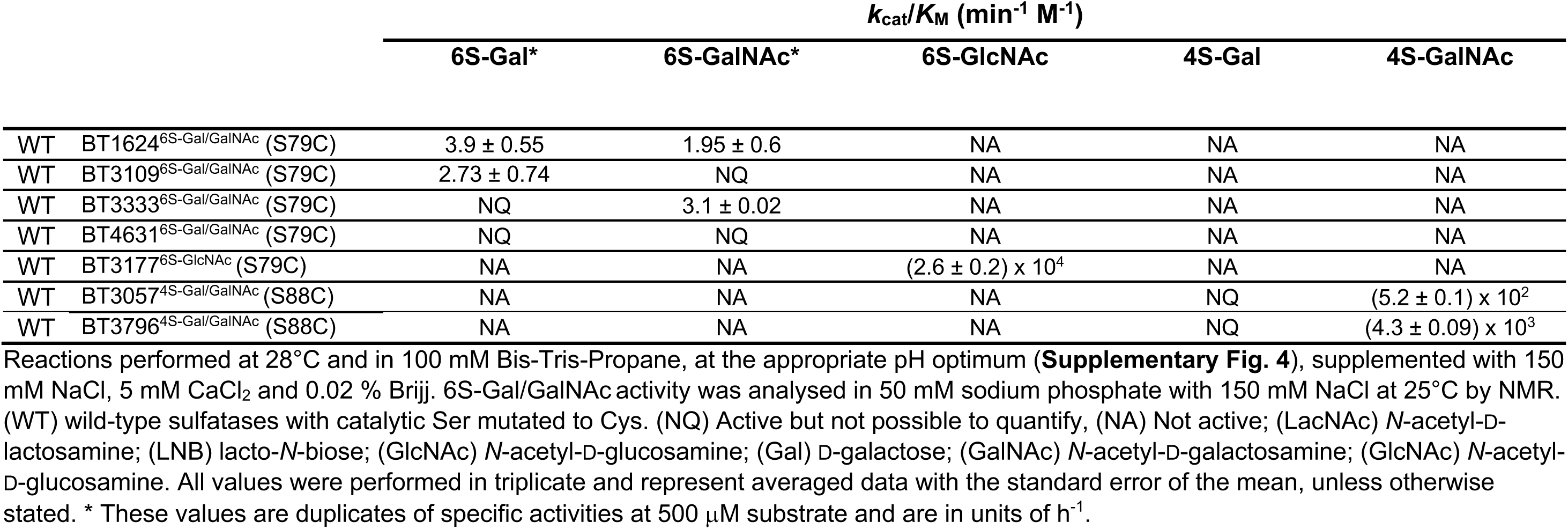
Sulfatase kinetics for WT and mutants against different saccharides

**Supplementary Table 4.** LC-MS analysis of colonic mucin oligosaccharides (cMO) Provided as a separate multi-tab Excel file

**Supplementary Table 5.** LC-MS analysis of O-glycans in culture supernatant of Δ*bt1636^3S-Gal^* mutant by LC-MS/MS Provided as a separate multi-tab Excel file

**Supplementary Table 6.**
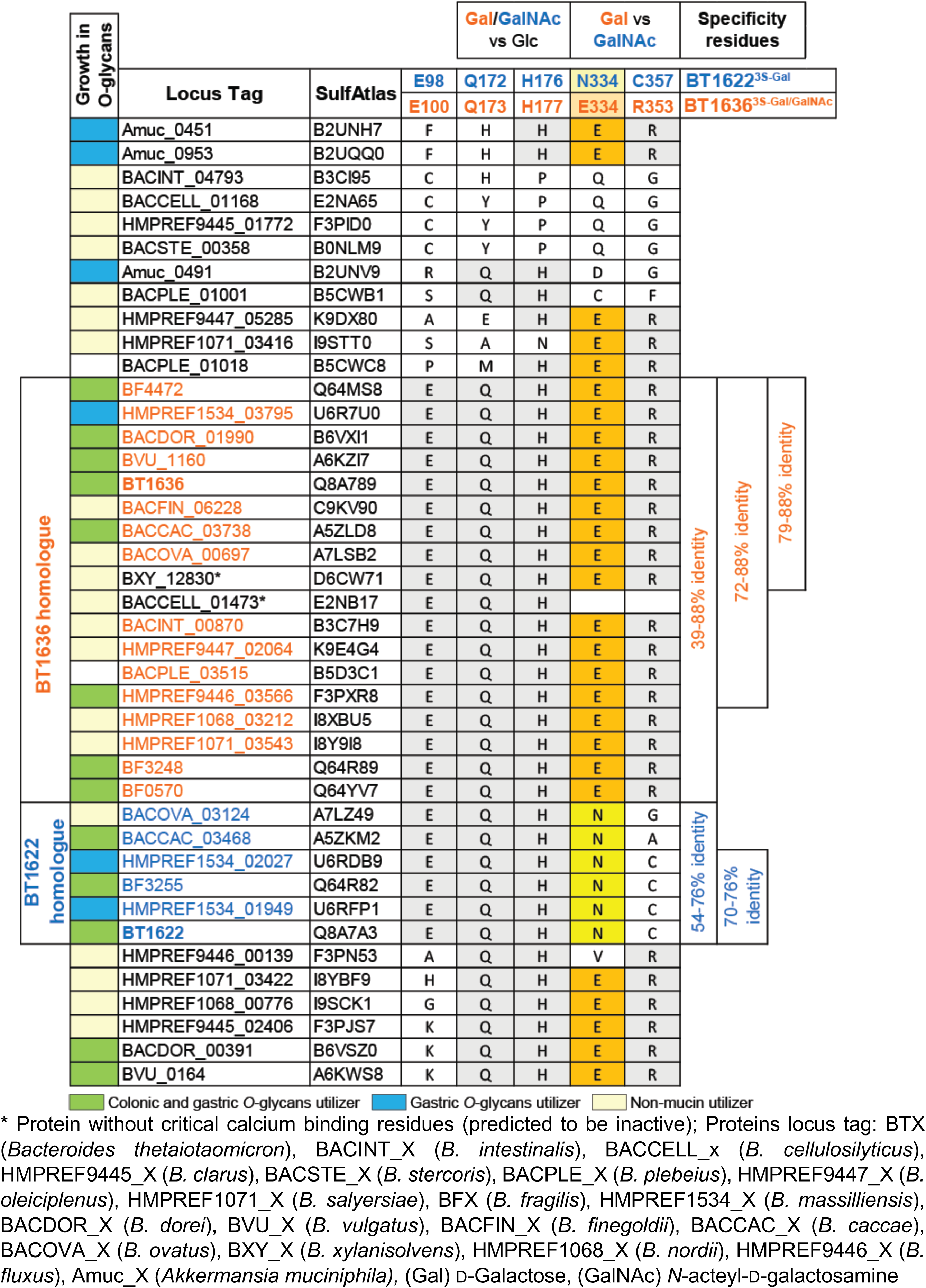
Conservation of S1_20 3S-Gal/GalNAc specificity residues in *Bacteroides* type strains and *Akkermansia muciniphila*

**Supplementary Table 7.**
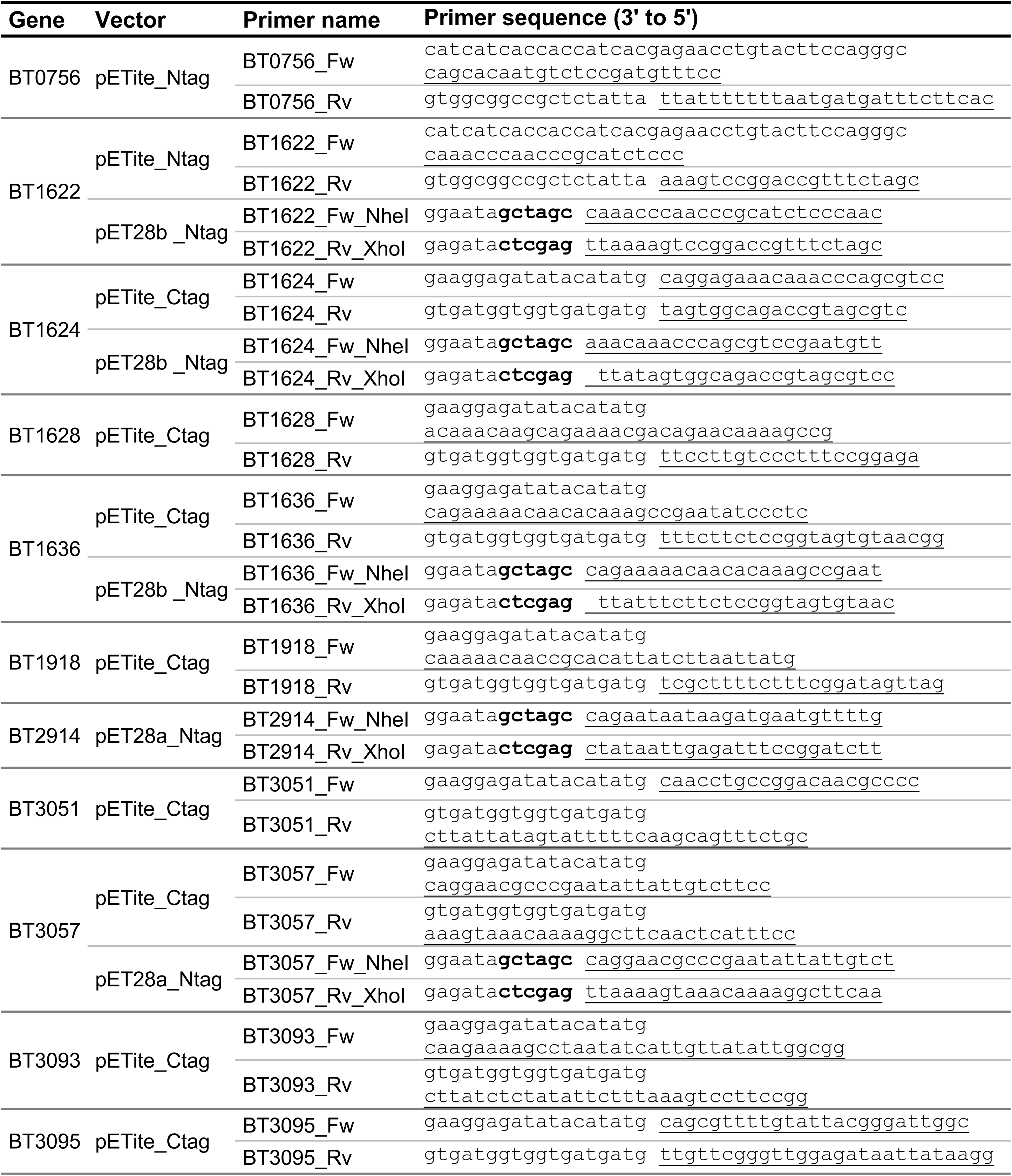
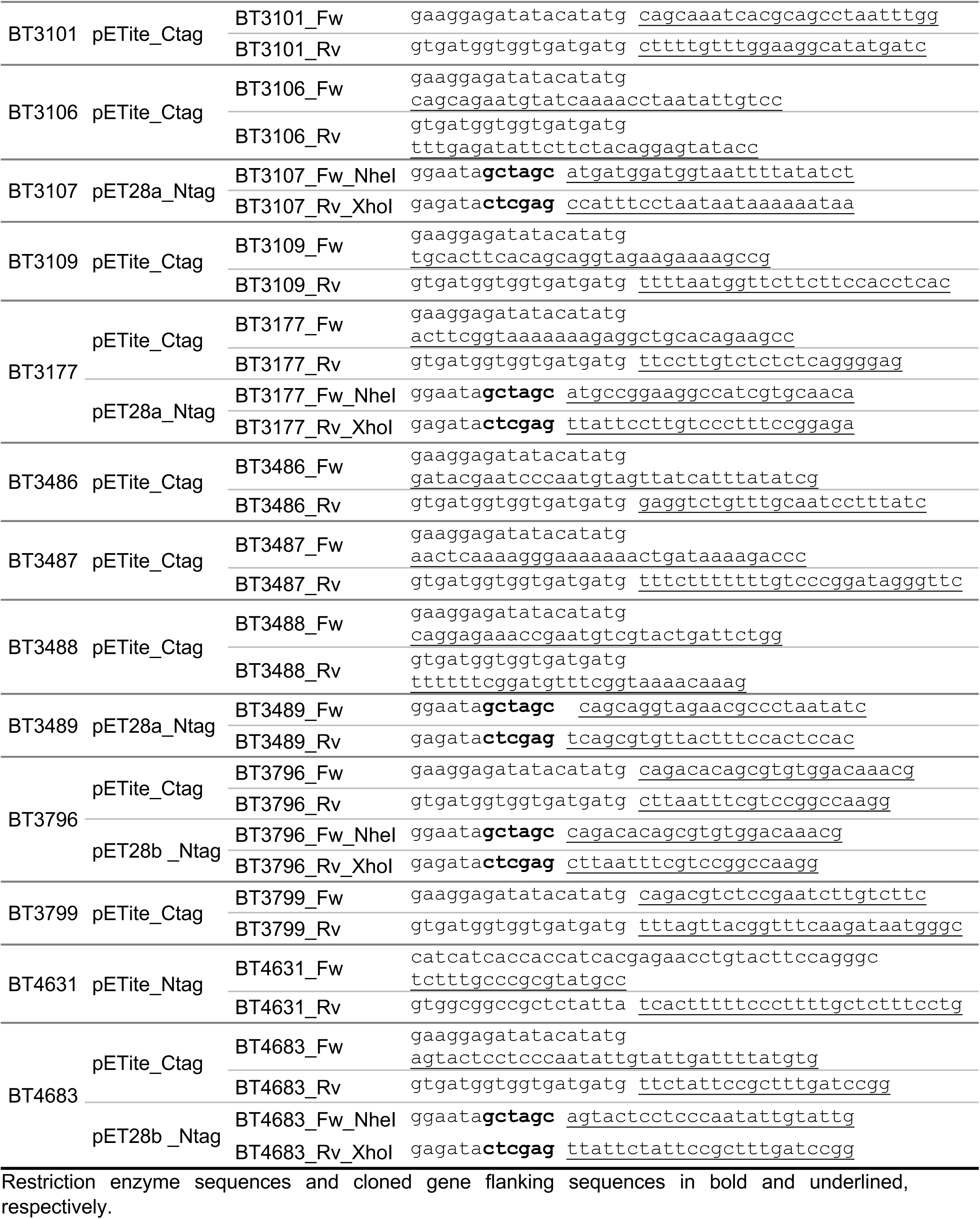
Primers designed to clone *Bt* sulfatases

**Supplementary Table 8.**
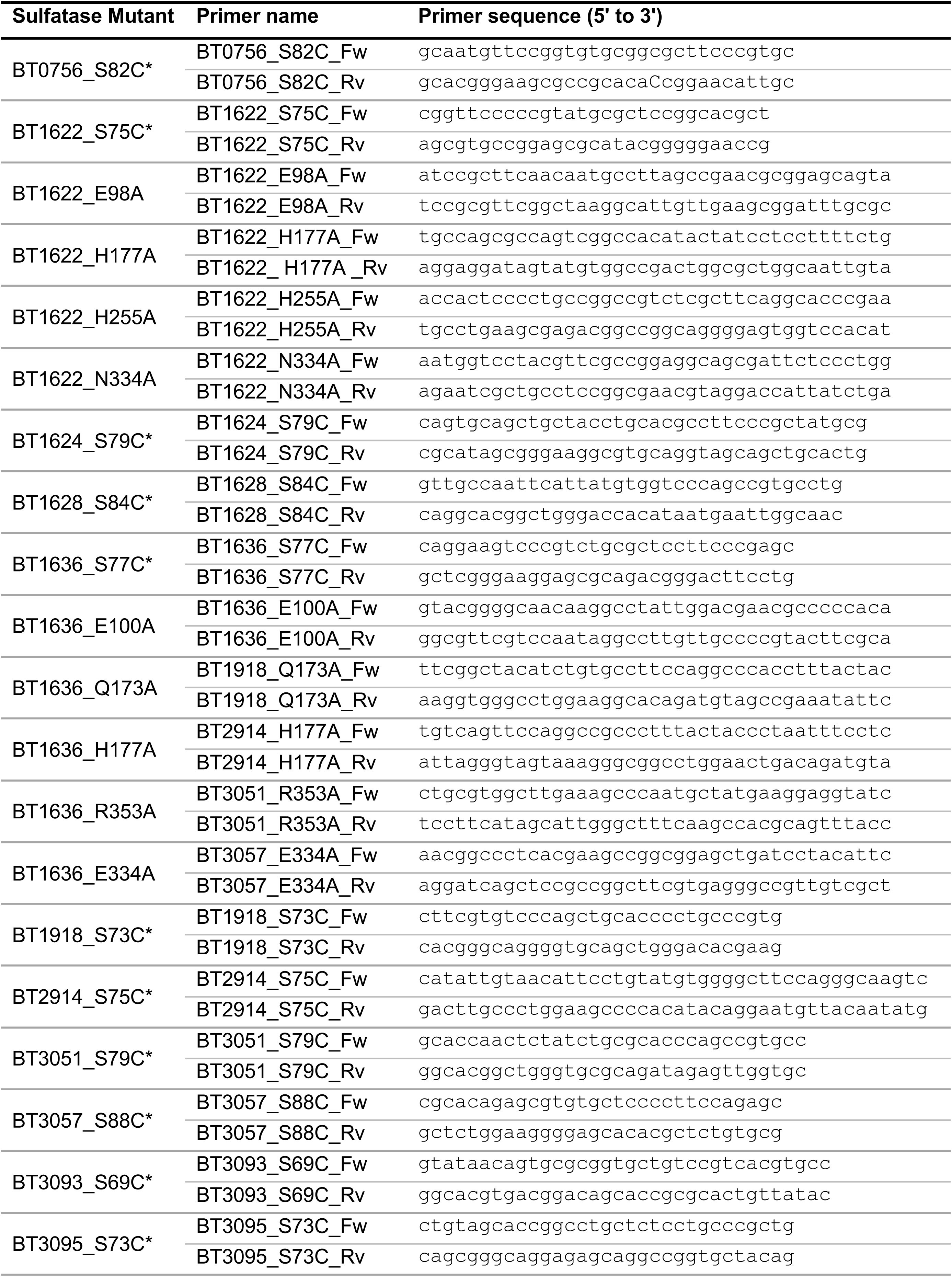
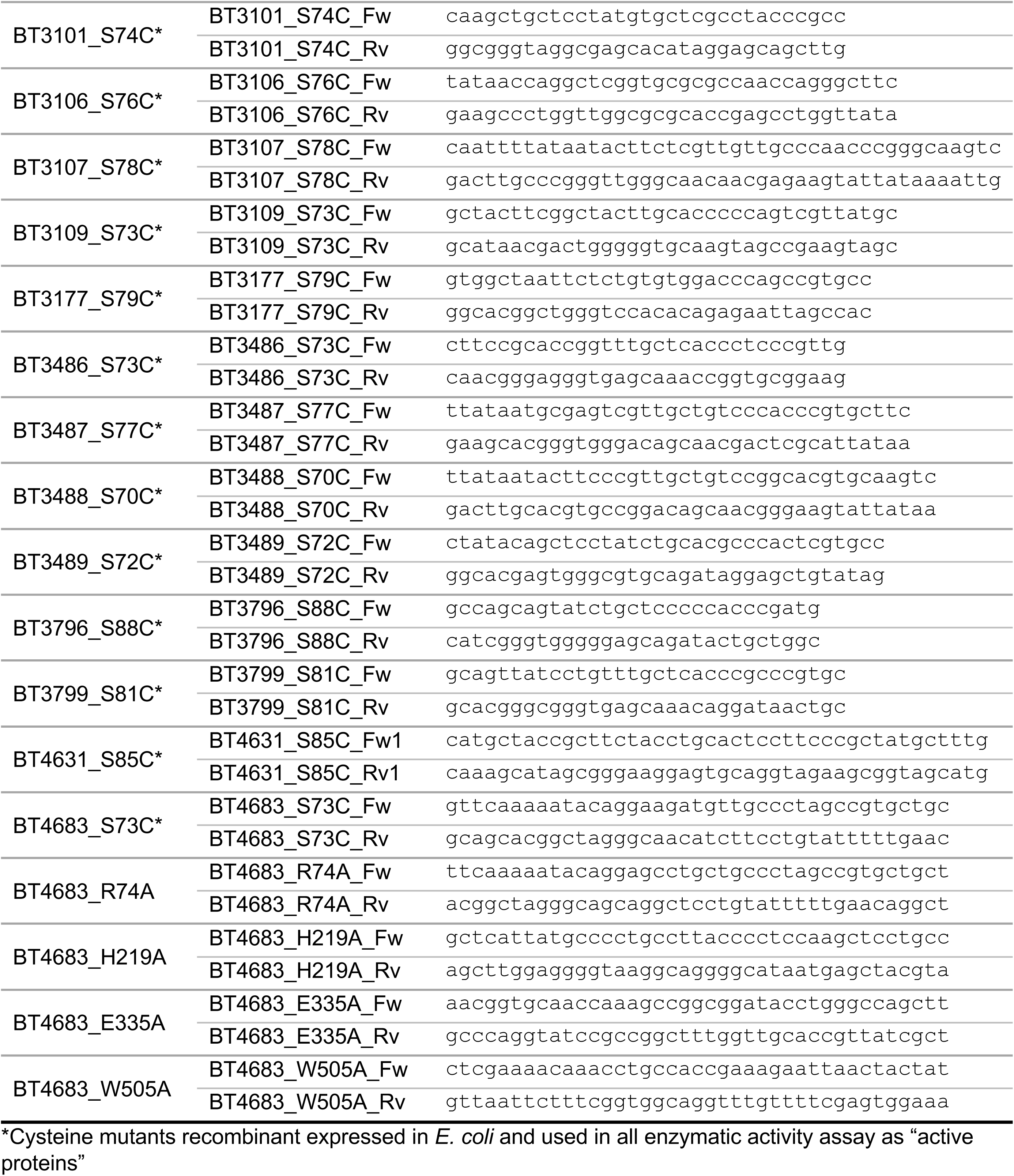
Primers designed to generate the site-directed mutants of *Bt* sulfatases

**Supplementary Table 9.**
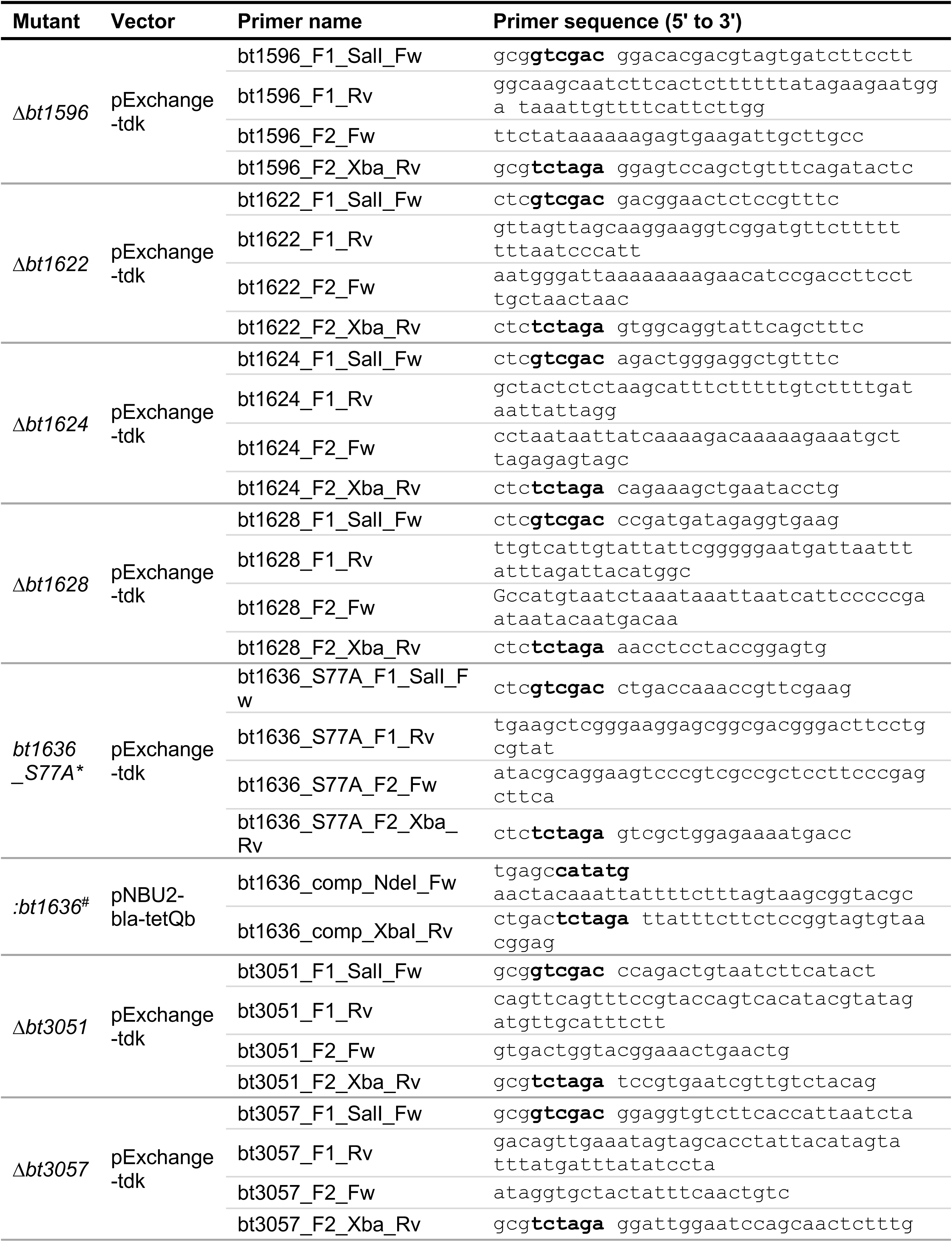
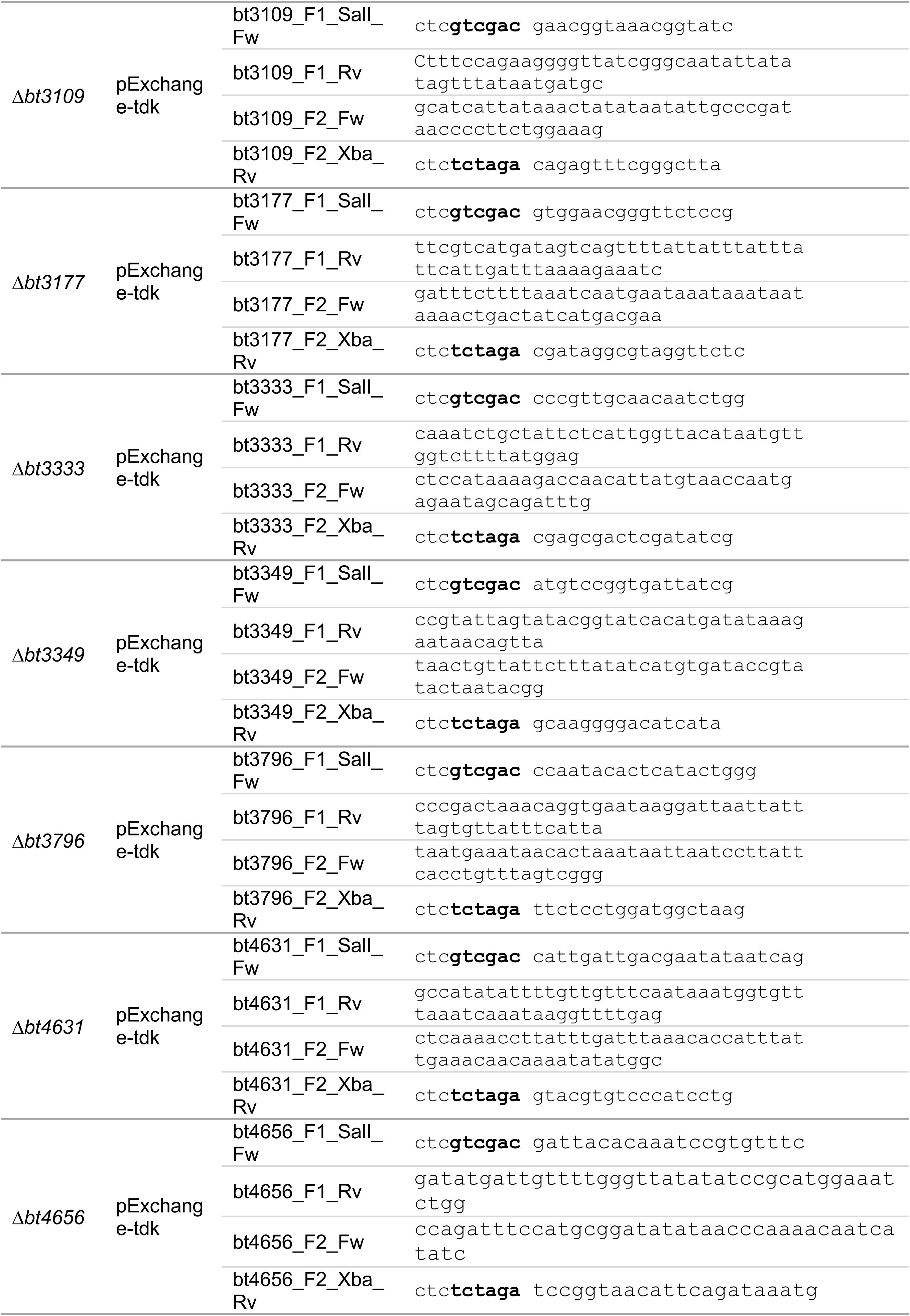
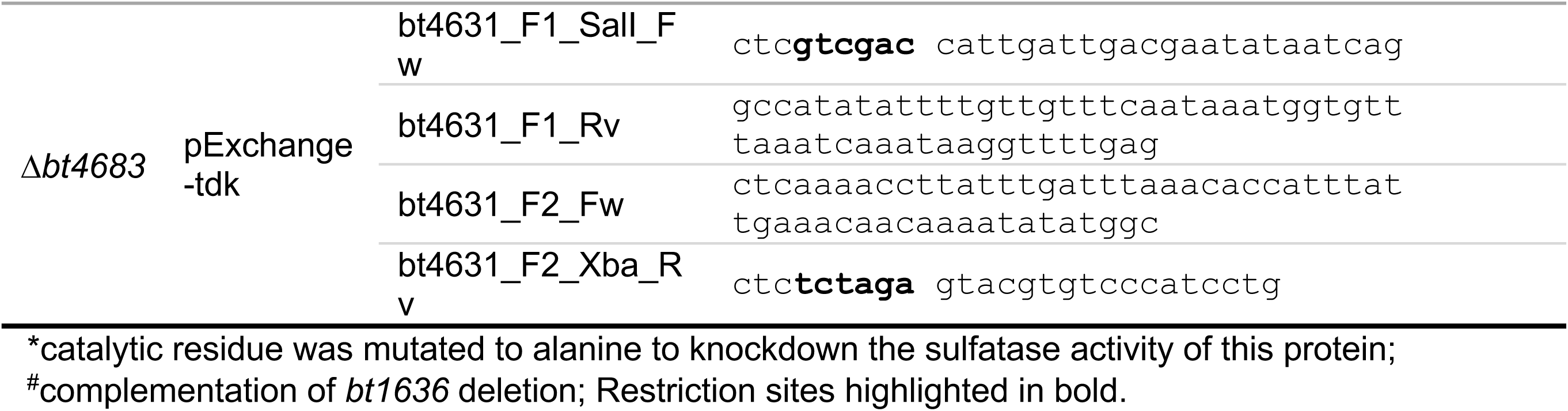
Primers designed to generate the in-frame gene deletions and complementations of *Bt* sulfatases

**Supplementary Table 10.**
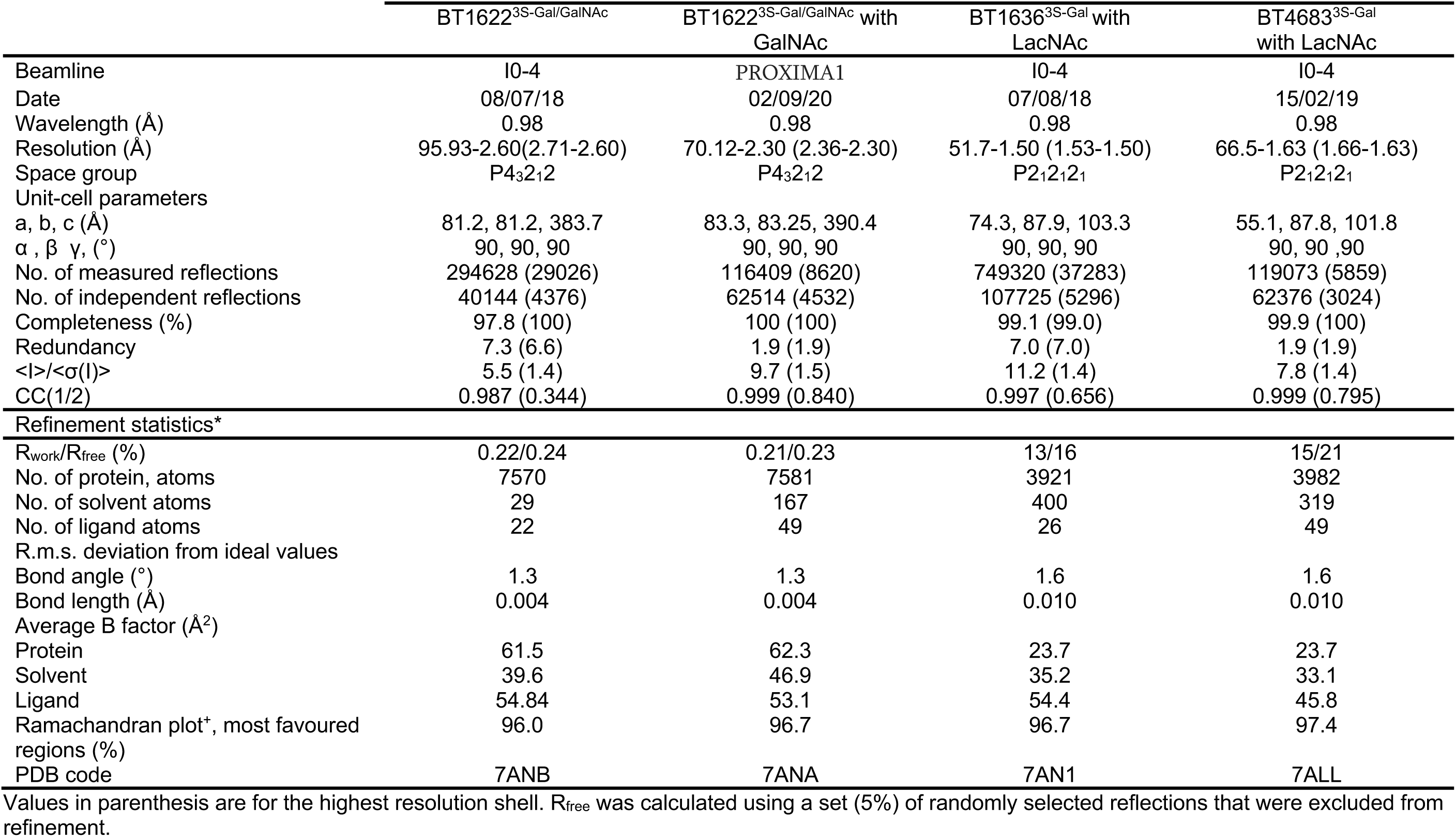

**Supplementary Table 11.**
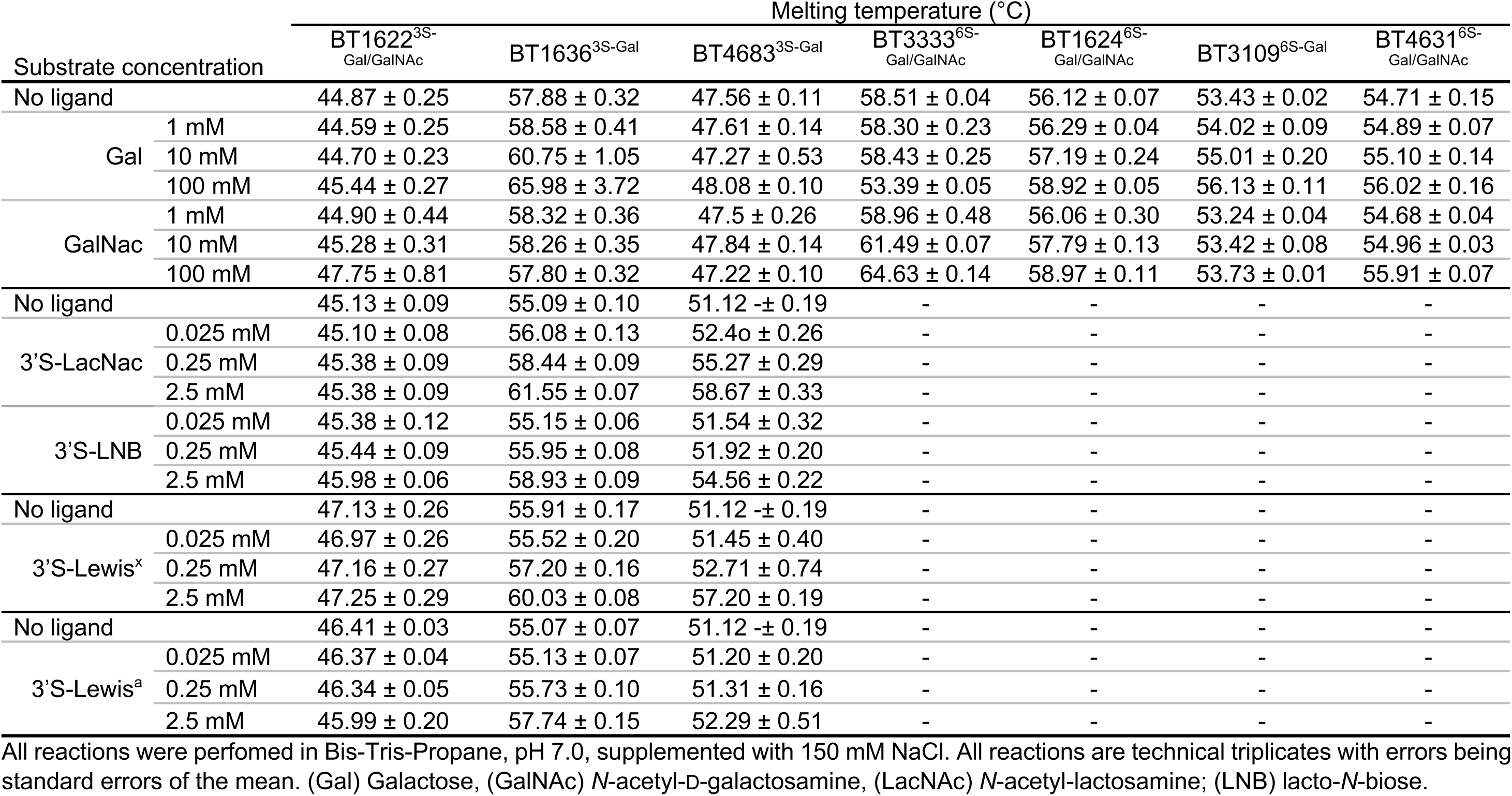
Melting temperatures of galactose targeting sulfatases with and without ligands.

**Supplementary Table 12.**
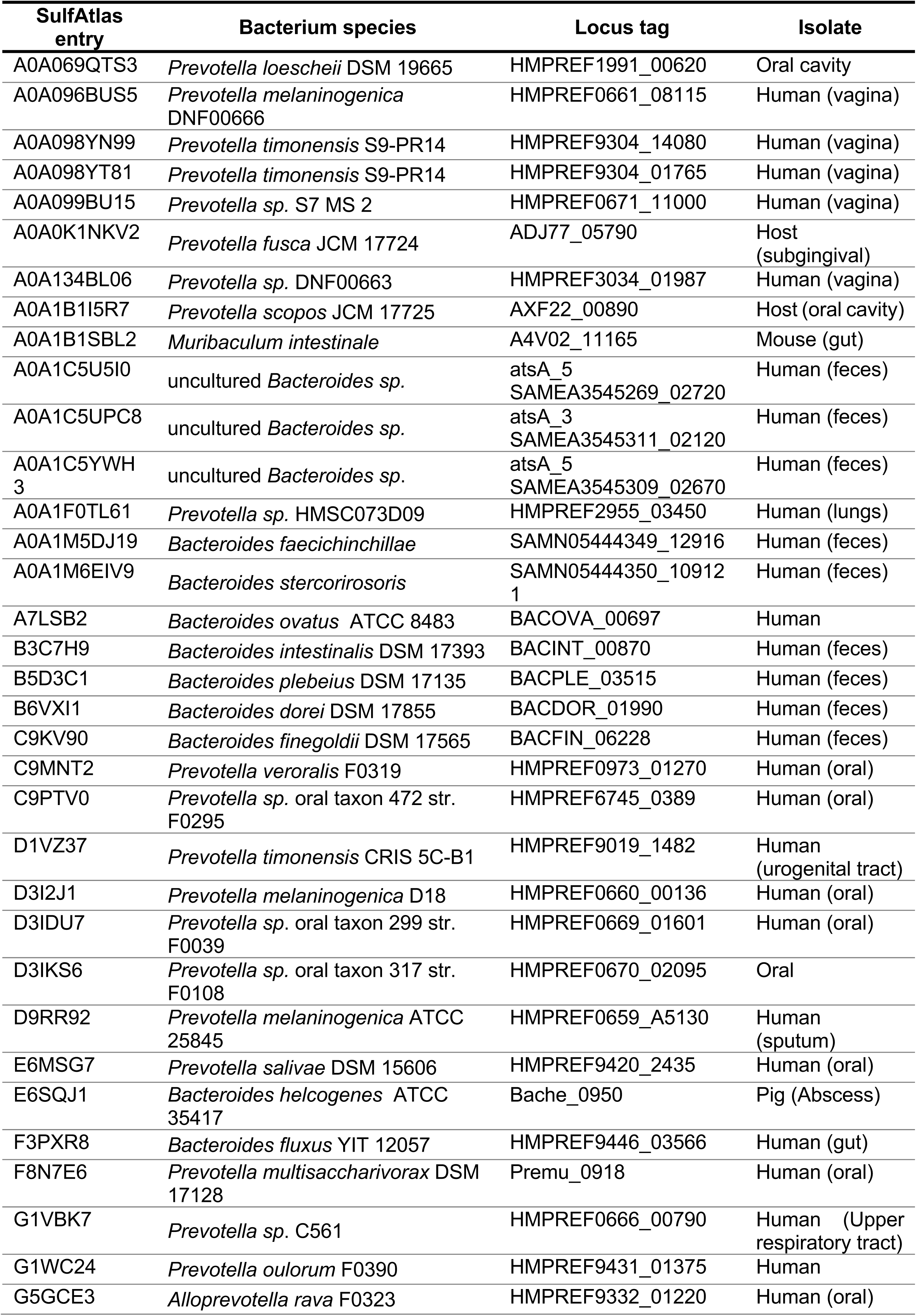
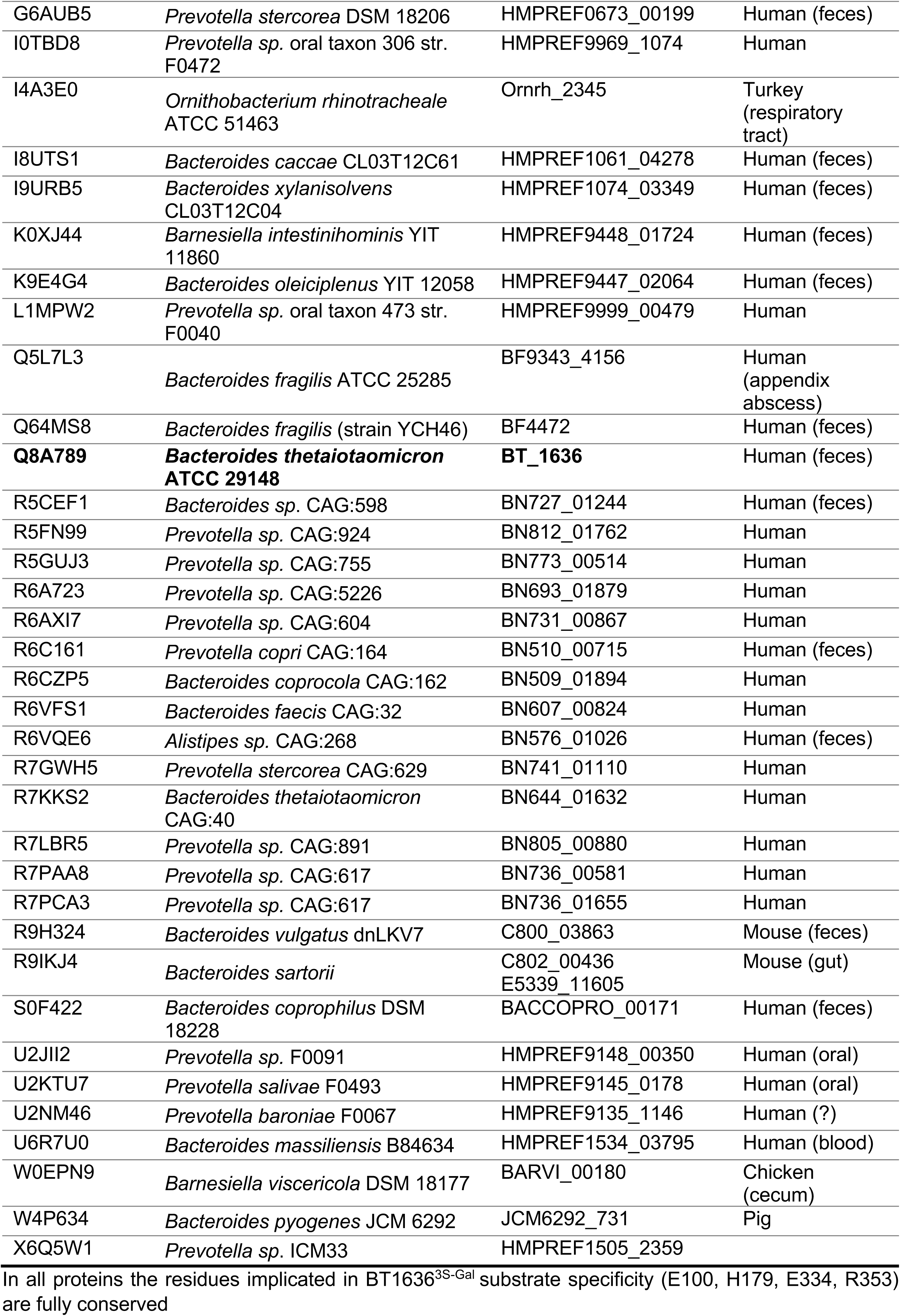
S1_20 homologues of BT1636^3S-Gal^

**Supplementary Table 13.**
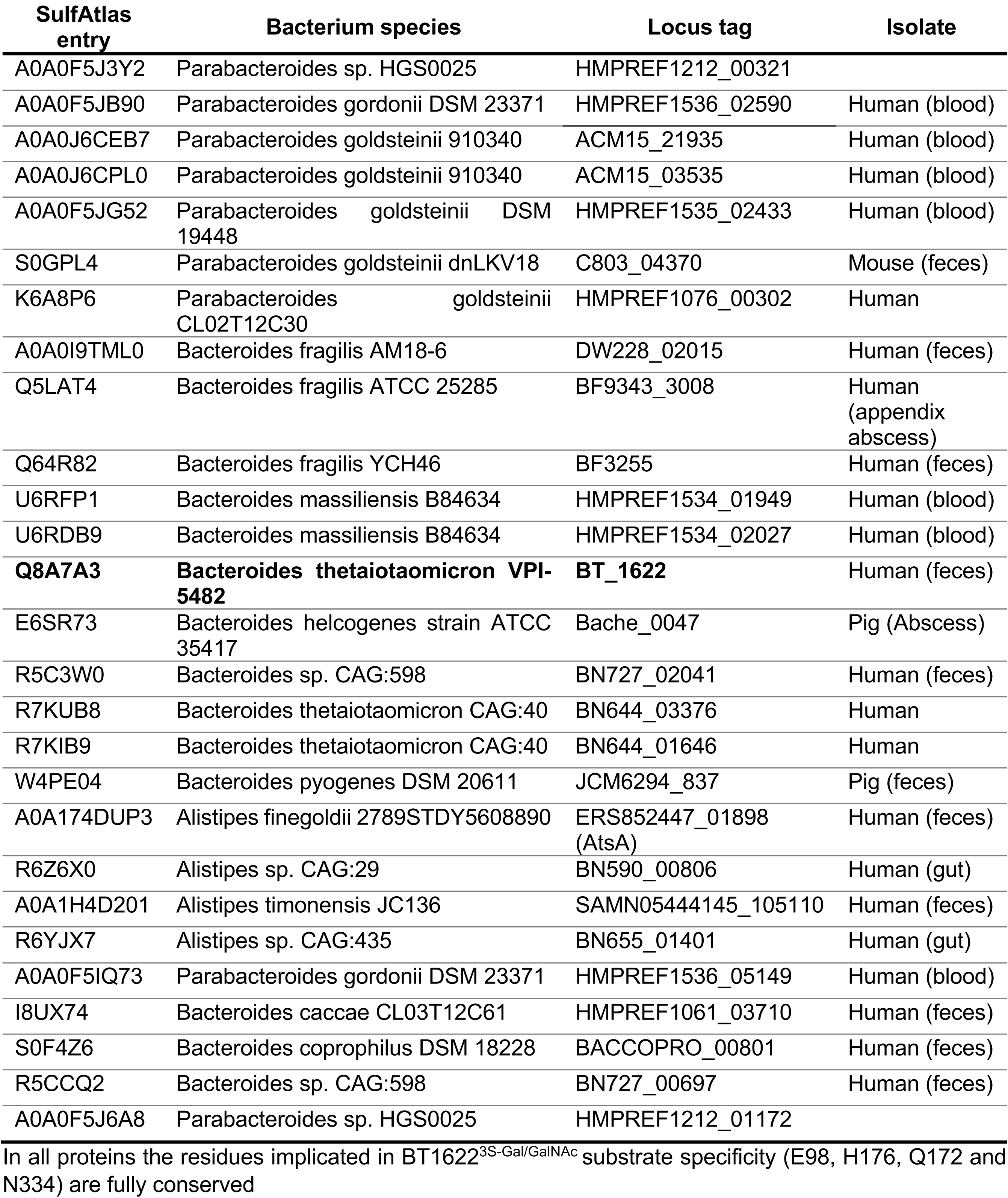
S1_20 homologues of BT1622^3S-Gal/GalNAc^

**Supplementary Figure 1.**
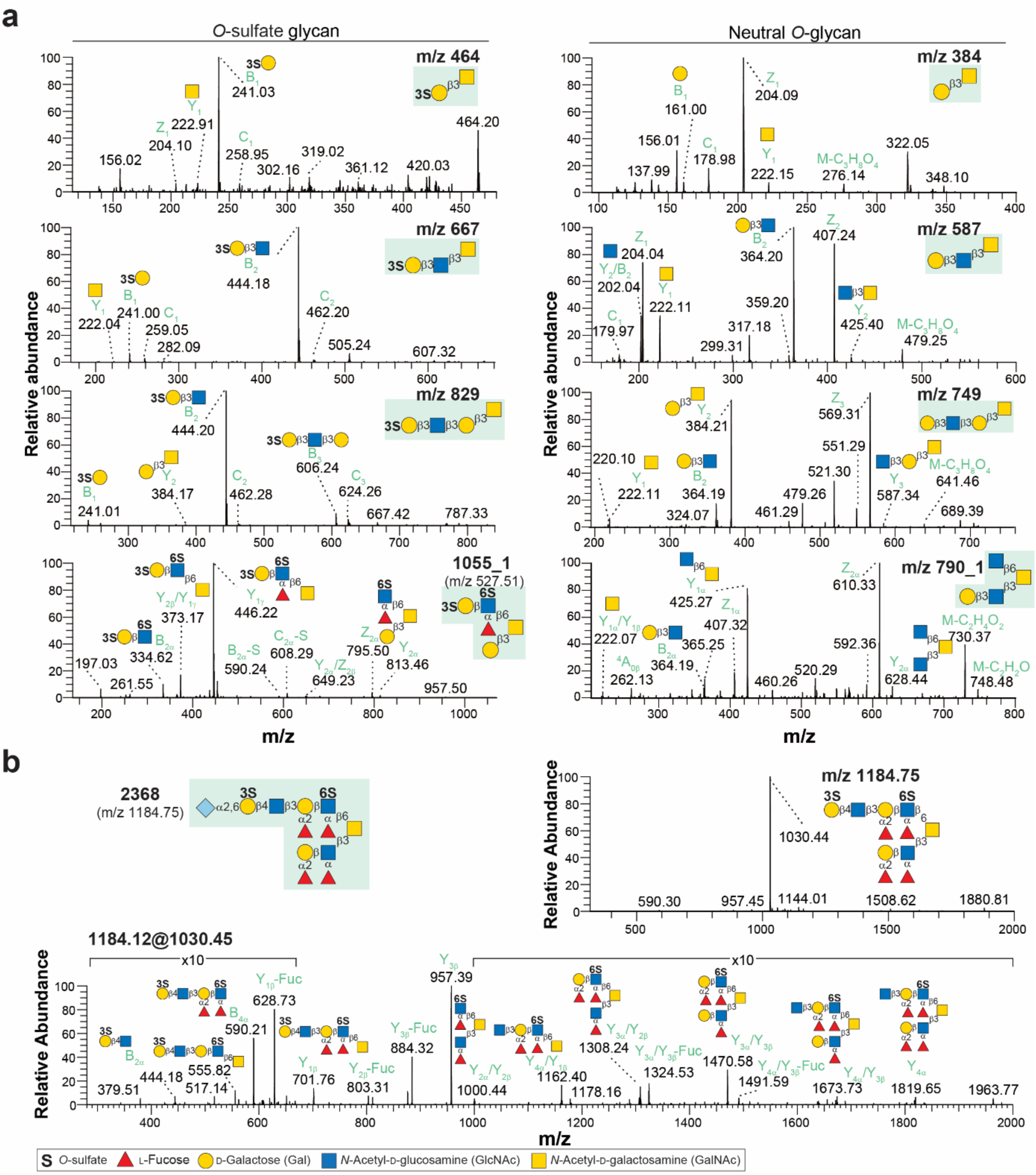
Characterization of negatively charged O-glycans from porcine colonic mucins using LC-MS/MS. **a,** MS/MS spectra of selected *O*-sulfate and neutral non-sulfated O-glycans glycan (left and right hand side panels, respectively). **b,** MS^2^ (m/z 1184.75, [M-2H]^2-^) and MS^3^ (m/z 1184.12@1030.45) spectra with a composition of NeuGc1Hex3HexNAc4deHex4Sul2.

**Supplementary Figure 2.**
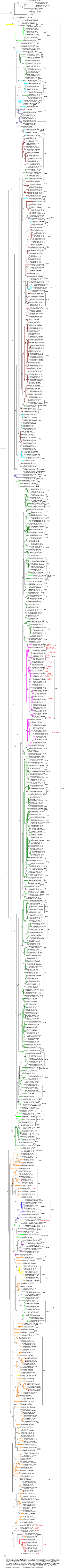
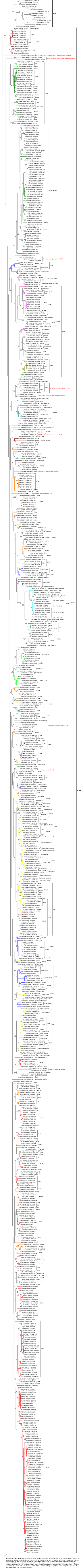
Phylogenetic analysis of S1_4 subfamily showing the conservation of BT4683^3S-Gal^ specificity residues Provided as separate PDF file due to size

**Supplementary Figure 3.** Phylogenetic analysis of S1_20 subfamily showing the conservation of BT1636^3S-Gal^ specificity residues Provided as separate PDF file due to size

**Supplementary Figure 4.**
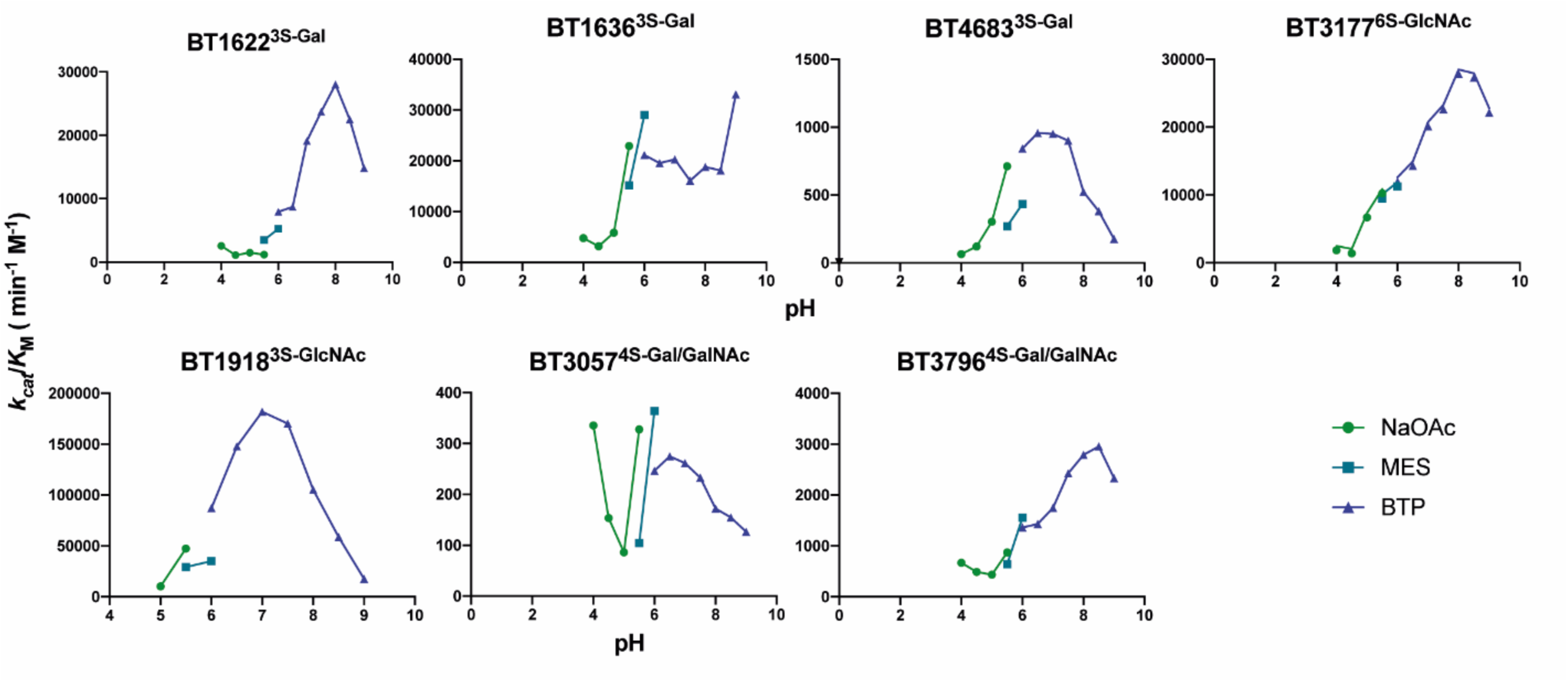
pH profile of different sulfatases. Graphics showing the pH optimum for BT1636^3S-Gal^ and BT3057^4S-Gal/GalNAc^ is 6; BT4683^3S-Gal^ had a pH optimum of 6.5; BT19183S-GlcNAc had a pH optimum of 7.0; BT1622^S-Gal^ and BT3177^6S-GlcNAc^ had a pH optimum of 8, whilst BT3796^4S-Gal/GalNAc^ had a pH optimum of 8.5. All reactions were performed using 100 mM of the appropriate buffer supplemented with 150 mM NaCl and 5 mM CaCl_2_. A substrate concentration of 1 μM was used and an enzyme concentration of between 0.1 – 2 μM was deployed depending on the enzyme.

